# Cigarette Smoke Activates NOTCH3 to Promote Goblet Cell Differentiation in Human Airway Epithelial Cells

**DOI:** 10.1101/2020.07.09.195818

**Authors:** Manish Bodas, Andrew R. Moore, Bharathiraja Subramaniyan, Constantin Georgescu, Jonathan D. Wren, Willard M. Freeman, Brent R. Brown, Jordan P. Metcalf, Matthew S. Walters

## Abstract

Chronic obstructive pulmonary disease (COPD) is the 3rd leading cause of death in the United States and is primarily caused by cigarette smoking. Increased numbers of mucus-producing secretory (“goblet”) cells defined as goblet cell metaplasia or hyperplasia (GCMH), contributes significantly to COPD pathophysiology. The objective of this study was to determine whether NOTCH signaling regulates goblet cell differentiation in response to cigarette smoke. Primary human bronchial epithelial cells (HBECs) from nonsmokers and COPD smokers were differentiated *in vitro* on air-liquid interface and exposed to cigarette smoke extract (CSE) for 7 days. NOTCH signaling activity was modulated using (1) the NOTCH/γ-secretase inhibitor Dibenzazepine (DBZ), (2) lentiviral over-expression of the NOTCH3-intracellular domain (NICD3) or (3) NOTCH3-specific siRNA. Cell differentiation and response to CSE were evaluated by qPCR, Western blotting, immunostaining and RNA-Seq. We found that CSE exposure of nonsmoker airway epithelium induced goblet cell differentiation characteristic of GCMH. Treatment with DBZ suppressed CSE-dependent induction of goblet cell differentiation. Furthermore, CSE induced NOTCH3 activation, as revealed by increased NOTCH3 nuclear localization and elevated NICD3 protein levels. Over-expression of NICD3 increased the expression of goblet cell associated genes SPDEF and MUC5AC, whereas NOTCH3 knockdown suppressed CSE-mediated induction of SPDEF and MUC5AC. Finally, CSE exposure of COPD airway epithelium induced goblet cell differentiation in a NOTCH3-dependent manner. These results identify NOTCH3 activation as one of the important mechanisms by which cigarette smoke induces goblet cell differentiation, thus providing a novel potential strategy to control GCMH-related pathologies in smokers and patients with COPD.

## Introduction

Chronic obstructive pulmonary disease (COPD) is characterized by a combination of pathological conditions including chronic bronchitis and emphysema (1, 2), and is the 3^rd^ leading cause of death in the United States (3). Exposure to cigarette smoke is the primary risk factor for development and progression of COPD (4). The earliest cigarette smoking-induced changes relevant to the pathogenesis of COPD occur in the airway epithelium (1, 2), a multi-cellular tissue that covers the luminal surface of the conducting airways from the proximally located trachea to the distal bronchioles (5, 6). This pseudostratified airway epithelium functions as a barrier to protect the lung from pathogens and harmful environmental factors via the action of specialized cells, including ciliated, secretory (goblet and club), basal and neuroendocrine cells (5–7). An intricate balance of the percentage of each cell type is critical to maintaining a healthy airway epithelium capable of efficient mucociliary clearance and air flow (6–9). Structural changes in airway epithelial architecture (termed “epithelial remodeling”) play a crucial role in COPD pathogenesis (1, 2). Goblet cell metaplasia or hyperplasia (GCMH) is an epithelial remodeling phenotype characterized by increased numbers of goblet cells (9, 10). These changes culminate in excess mucus production that impairs mucociliary clearance, increases airway resistance, and contributes significantly to COPD morbidity and mortality (1, 9). Therefore, identifying the mechanisms by which cigarette smoke exposure regulates goblet cell differentiation is critical to developing novel therapeutics to treat COPD-associated GCMH.

One such potential mechanism is the NOTCH signaling pathway, which plays a vital role in regulating differentiation of the airway epithelium in both humans and mice (11), including goblet cell differentiation (12–22). NOTCH signaling is an evolutionary conserved pathway that functions in a cell-cell contact-dependent manner. Activation of canonical NOTCH signaling involves the binding of a plasma membrane-bound ligand (DLL1, DLL3, DLL4, JAG1 or JAG2) to one of the four receptors (NOTCH1-4) located on the plasma membrane of a neighboring cell (23). Upon ligand binding, the receptor is activated by multiple cleavage events, the final of which is mediated by the γ-secretase enzyme complex that results in release of the Notch intracellular domain (NICD) into the cytoplasm. The NICD subsequently translocates to the nucleus and induces transcription of multiple target genes (23). Previous studies have demonstrated the role of NOTCH signaling in the development of GCMH using models involving exposure to allergens, inflammatory cytokines and viral infections (12, 15, 17–19), but whether NOTCH is involved in cigarette smoke induced GCMH is unexplored.

In the present study, we hypothesized that cigarette smoke exposure increases NOTCH signaling activation in the airway epithelium leading to increased goblet cell differentiation and development of GCMH. To investigate this, we exposed air-liquid interface (ALI) cultures of primary human bronchial epithelial cells (HBECs) from nonsmokers and COPD patients with cigarette smoke extract (CSE). Our data demonstrate that CSE exposure of the airway epithelium generated from nonsmoker and COPD HBECs induces goblet cell differentiation and increases the numbers of MUC5AC^+^ goblet cells characteristic of GCMH. CSE exposure induced activation of NOTCH3 signaling and over-expression of the constitutively active NICD3 increased expression of the goblet cell associated genes SPDEF and MUC5AC in the absence of CSE. Finally, siRNA mediated knockdown of NOTCH3 in nonsmoker and COPD HBECs suppressed CSE-mediated induction of SPDEF and MUC5AC. Therefore, our results identify NOTCH3 signaling as one of the mechanisms by which cigarette smoke exposure induces goblet cell differentiation in the airway epithelium. This information is critical to the understanding of the pathophysiology of COPD and provides a potential strategy to control GCMH-related pathologies in smokers and patients with COPD.

## Methods

Full details of the methods are provided in the online data supplement.

### Primary human bronchial epithelial cell (HBEC) culture

Primary HBECs were purchased (Lonza, Morristown, NJ, USA) from nonsmokers (Cat# CC-2540) and COPD smokers (Cat# 00195275 and 00195275S). The demographic details of the donors is summarized in Table E1. Cells were maintained in BronchiaLife™ epithelial airway medium (BLEAM) (Cat# LL-0023, Lifeline® Cell Technology, Oceanside, CA, USA) supplemented with Penicillin (100 I.U./ml) - Streptomycin (100 μg/ml) as previously described (14).

### Air-liquid interface (ALI) culture

Briefly, 1 × 10^5^ HBECs in 100 μl of BLEAM media were seeded in the apical chamber of a Transwell® insert (Cat# 3470, Corning^®^, Corning, NY, USA) pre-coated with human type IV collagen (Cat# C7521, Sigma Aldrich, St. Louis, MO, USA) with 1 ml of BLEAM in the basolateral chamber (ALI day −2). The following day, fresh BLEAM media was replaced in the apical and basolateral chambers (100 μl and 1 ml, respectively). Following two days of submerged culture, media from the apical chamber was removed to expose the cells to air (ALI day 0), and 1 ml of HBTEC ALI differentiation medium (Cat# LM-0050, Lifeline^®^ Cell Technology) supplemented with Penicillin (100 I.U./ml) - Streptomycin (100 μg/ml) was added to the basolateral chamber. The media was replaced every other day.

### Cigarette smoke extract (CSE) and DBZ treatments

Cells were treated with CSE (up to 2.5%) via the basolateral chamber for 7 days (either ALI day 0-7 or ALI day 28-35) with freshly thawed CSE added at each media change. For experiments involving suppression of NOTCH signaling, cells were treated with CSE and/or DBZ (0.1 μM, Cat# 565789, EMD Chemicals Inc., Gibbstown, NJ, USA) from ALI day 28 to 35, with an equal volume of DMSO (Sigma Aldrich) used as vehicle control.

### RNA sequencing

RNA sequencing was performed on a NextSeq 500 Flowcell, High SR75 (Illumina, San Diego, CA, USA) following library preparation using the QuantSeq 3′ mRNA-Seq Library Prep Kit FWD for Illumina (Lexogen, Vienna, Austria). The raw data are publicly available at the Gene Expression Omnibus (GEO) site (http://www.ncbi.nlm.nih.gov/geo/), accession number GSE152446. Genes that were differentially expressed in response to CSE treatment were determined using a threshold of p<0.05 on the false discovery rate (FDR).

### Lentivirus-based overexpression of NICD3

Generation of control or NICD3 expressing replication deficient lentiviruses was described previously (14).

### siRNA mediated knockdown of NOTCH3

HBECs were either transfected with 1 pmol of control siRNA (Cat# 4390844, Thermo Scientific) or NOTCH3 siRNA (Cat # 4392420, Thermo Scientific, Waltham) using Lipofectamine RNAiMax Reagent (Cat# 13778-045, Thermo Scientific) and OptiMEM media (Cat# 1985-010, Thermo Scientific) at the time of seeding the cells in ALI culture.

### Statistics

Statistical analysis was performed by two-tailed Mann Whitney U test using the GraphPad Prism version 8.0. A p-value of ≤ 0.05 was considered significant.

## Results

### In vitro cigarette smoke extract exposure promotes goblet cell differentiation

To study how cigarette smoke exposure induces goblet cell differentiation in the airway epithelium, primary HBECs from nonsmokers were cultured *in vitro* at ALI for 28 days and then exposed to different concentrations of CSE for 7 days (Figure 1A). Expression of the oxidative-stress response gene CYP1A1 was significantly upregulated in the presence of 0.5% (10.8 fold) and 2.5% (92.6 fold) CSE in a concentration-dependent manner confirming that the cells were responding to CSE exposure (Figure 1B). H&E staining of paraffin-embedded sections demonstrated that exposure to CSE led to the development of epithelial remodeling with the appearance of a disordered, thickened epithelium compared to untreated cells (Figure 1C). Analysis of cell-type specific marker expression showed no significant change in expression of basal (KRT5) and ciliated (DNAI1) cell markers in CSE (both 0.5% and 2.5%) treated cells compared to untreated controls (Figure 1D). However, CSE treatment significantly downregulated expression of the club cell marker SCGB1A1 (0.59 fold, 0.5% CSE and 0.47 fold, 2.5% CSE) and significantly increased expression of the goblet cell marker MUC5AC (10.4 fold, 0.5% CSE and 19.1 fold, 2.5% CSE) (Figure 1D). To validate that the gene expression changes observed for the cell-type specific markers reflected alterations in cell populations, we quantified the number of basal, ciliated, club and goblet cells histologically (Figure 1E). Similar to the gene expression data, there were no significant changes in the numbers of basal (KRT5^+^) and ciliated (acetylated tubulin^+^) cells in CSE treated cells compared to untreated controls, as observed by immunostaining. Furthermore, a significant decrease in club cell numbers (SCGB1A1^+^) was observed for both 0.5% (0.51 fold) and 2.5% (0.45 fold) CSE treatment. However, only treatment with 2.5% CSE resulted in a significant increase in the numbers of MUC5AC^+^ goblet cells (4.3 fold). These data demonstrate that treatment of *in vitro* differentiated airway epithelium with 2.5% CSE promotes goblet cell differentiation and induces epithelial remodeling characteristic of GCMH.

**Figure 1.**
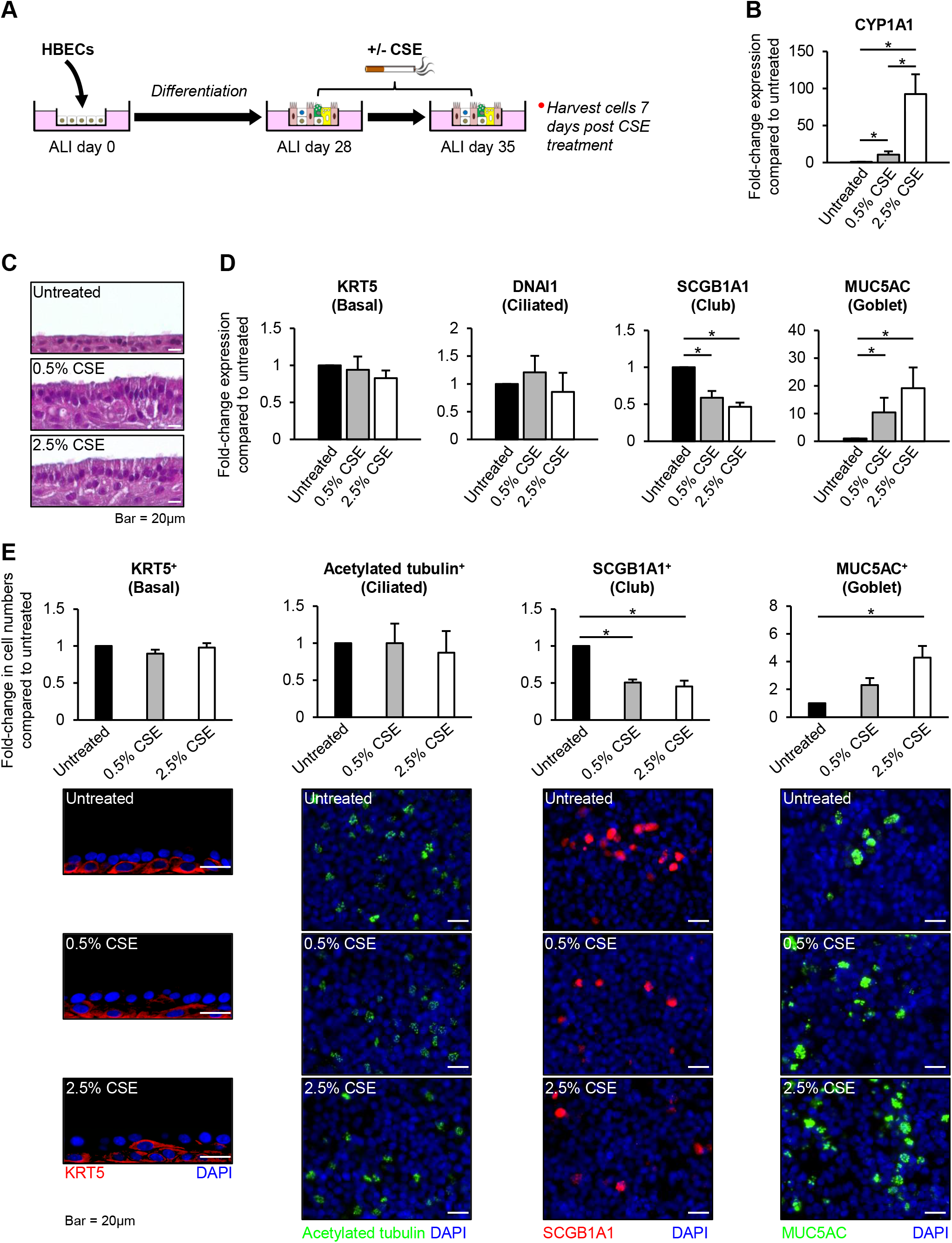
*In vitro* CSE exposure of nonsmoker airway epithelium. **(A)** Schematic of air-liquid interface (ALI) model. Primary human bronchial epithelial cells (HBECs) from nonsmokers were cultured on ALI for 28 days to differentiate into a pseudostratified epithelium containing basal, ciliated and secretory (club and goblet) cells. At ALI day 28, the cultures are then untreated or treated for 7 days with cigarette smoke extract (CSE). **(B)** qPCR of CYP1A1 expression. Data represented as mean fold-change in expression compared to untreated cells from n=4 independent experiments. **(C)** H&E staining, Scale bar = 20μm. **(D)** qPCR of cell type specific marker expression. Data presented as mean fold-change in expression compared to untreated cells from n=4 independent experiments. **(E)** Immunofluorescent staining of basal cells (KRT5, red), ciliated cells (acetylated tubulin, green), club cells (SCGB1A1,), and goblet cells (MUC5AC, green). Images were quantified using the ImageJ software (NIH). Data presented as mean fold-change in cell numbers compared to untreated cells from n=4 independent experiments (see Figure E1 for presentation of these data as percentage positive cells for each condition). Scale bar = 20μm. For panels **B**, **D** and **E** the error bars indicate SEM. * p<0.05.

### Genome-wide transcriptome changes in response to CSE treatment

To identify candidate genes and/or pathways that regulate goblet cell differentiation in response to CSE, RNA-Seq was performed on untreated and 2.5% CSE treated airway epithelium, 7 days post CSE treatment (ALI day 35). Comparison of CSE-treated *vs.* untreated cells identified 273 genes (124 up-regulated and 149 down-regulated) with significant (FDR adjusted p<0.05) expression changes (File E1). Ingenuity Pathway Analysis demonstrated significant enrichment of molecular pathways previously associated with cigarette smoking and smoking-induced lung disease including HIF1α signaling (10 genes), nicotine degradation II (6 genes), and aryl hydrocarbon receptor signaling (9 genes) (Figure 2A and File E2) (24, 25). Furthermore, CSE exposure led to expression changes in many genes associated with goblet cell biology (9, 26–29) including inflammatory mediators (CXCL3, CXCL5, IL1β, IL1RN and IL19), growth factors and signal transduction (AREG, S100A8, S100A9 and S100A12), endoplasmic reticulum stress (CREB3L1, and ERN2), mucus hypersecretion (AQP5, ATP12A, CLCA2, SLC12A2, SLC26A4 and SLC31A1) and NOTCH signaling (JAG2, NOTCH1 and SPDEF) (Figure 2B and File E1).

**Figure 2.**
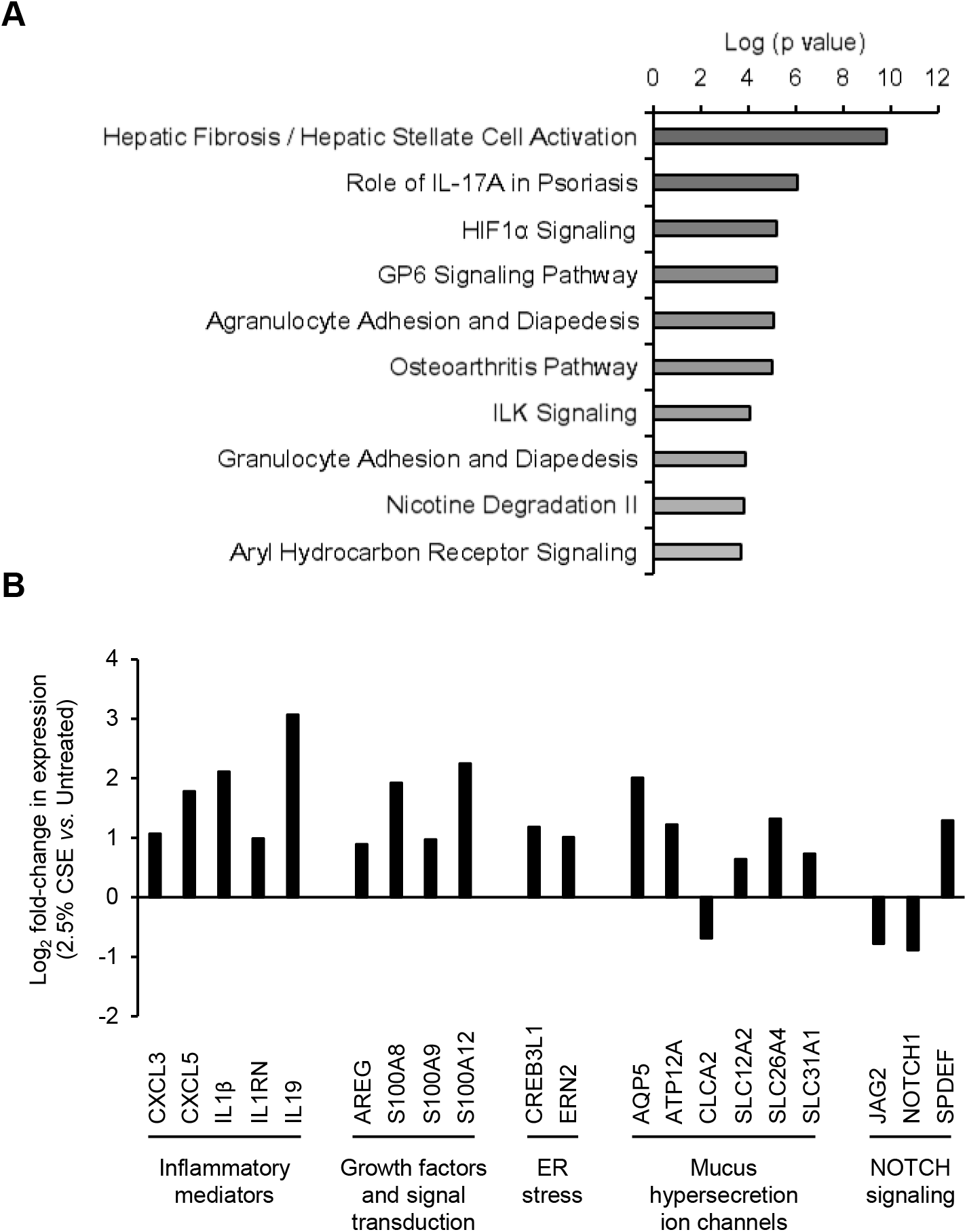
RNA-seq analysis to identify CSE-dependent transcriptome changes. Primary HBECs from nonsmokers were cultured on air-liquid interface (ALI) for 28 days to differentiate into a pseudostratified epithelium containing basal, ciliated and secretory (club and goblet) cells. At ALI day 28 the cultures were untreated or treated with 2.5% CSE for 7 days then harvested for RNA-Seq analysis (n=4 independent experiments). **(A)** Pathways enriched in the 273 CSE-responsive gene list on the basis of Ingenuity Pathway Analysis (IPA). Shown are the top ten IPA-enriched pathways based on P value (log-transformed). **(B)** Expression changes in genes associated with goblet cell biology present in the 273 CSE-responsive gene list. Data presented as mean Log2 fold-change in expression (2.5% CSE treated *vs.* Untreated) from n=4 independent experiments.

### NOTCH signaling activation regulates CSE-dependent goblet cell differentiation

NOTCH signaling (12–22) and SPDEF (30) play a critical role in regulation of airway epithelial goblet cell differentiation, with recent studies demonstrating that SPDEF expression is positively regulated by NOTCH signaling (12). Moreover, our RNA-Seq data showed that CSE exposure leads to increased expression of SPDEF suggestive of increased NOTCH signaling activity (Figure 2B). To ask whether global inhibition of NOTCH signaling by DBZ treatment can suppress CSE-mediated induction of goblet cell differentiation, primary HBECs from nonsmokers were cultured on ALI for 28 days and then exposed to CSE in the absence or presence of DBZ (Figure 3A). As controls, cells were treated with either DMSO (vehicle control) or DBZ in the absence of CSE. Morphological analysis (via H&E staining) of paraffin-embedded sections demonstrated that CSE exposure induced epithelial remodeling with the appearance of a thickened epithelium (Figure 3B). However, treatment with CSE+DBZ resulted in the appearance of an epithelium similar to DMSO or DBZ treated cells, suggesting that CSE-induced changes are regulated in a NOTCH signaling dependent manner. To further investigate the morphological changes in response to CSE +/- DBZ treatment, we quantified the number of basal, ciliated, club and goblet cells histologically (Figure 3C-F). Consistent with previous studies demonstrating a role for NOTCH signaling in regulating the balance of ciliated *vs.* secretory cell differentiation in adult airway epithelium (14, 15, 18–20, 31, 32), DBZ treatment alone led to a significant increase (2.01 fold) in ciliated (acetylated tubulin^+^) cell numbers and a significant decrease in the numbers of both SCGB1A1^+^ club cells (0.16 fold) and MUC5AC^+^ goblet cells (0.61 fold) compared to DMSO treated controls (Figure 3C-F). However, no significant changes in basal (KRT5^+^) cell numbers were observed (Figure 3C). Furthermore, as compared to DMSO treated cells, CSE treatment alone led to no significant changes in basal or ciliated cell numbers (Figure 3C-D). However, CSE treatment significantly decreased (0.46 fold) the numbers of club cells and significantly increased (3.32 fold) the numbers of goblet cells (Figure 3E and F) as expected. Additionally, a comparison of CSE treated *vs* CSE+DBZ treated cells demonstrated no significant changes in the numbers of basal or ciliated cells (Figure 3C-D), however a significant decrease in club cells and goblet cells was observed in CSE+DBZ treated cells (Figure 3E-F). The ability of DBZ to suppress the CSE-mediated increase in goblet cell numbers suggests that CSE promotes goblet cell differentiation via a NOTCH-dependent mechanism.

**Figure 3.**
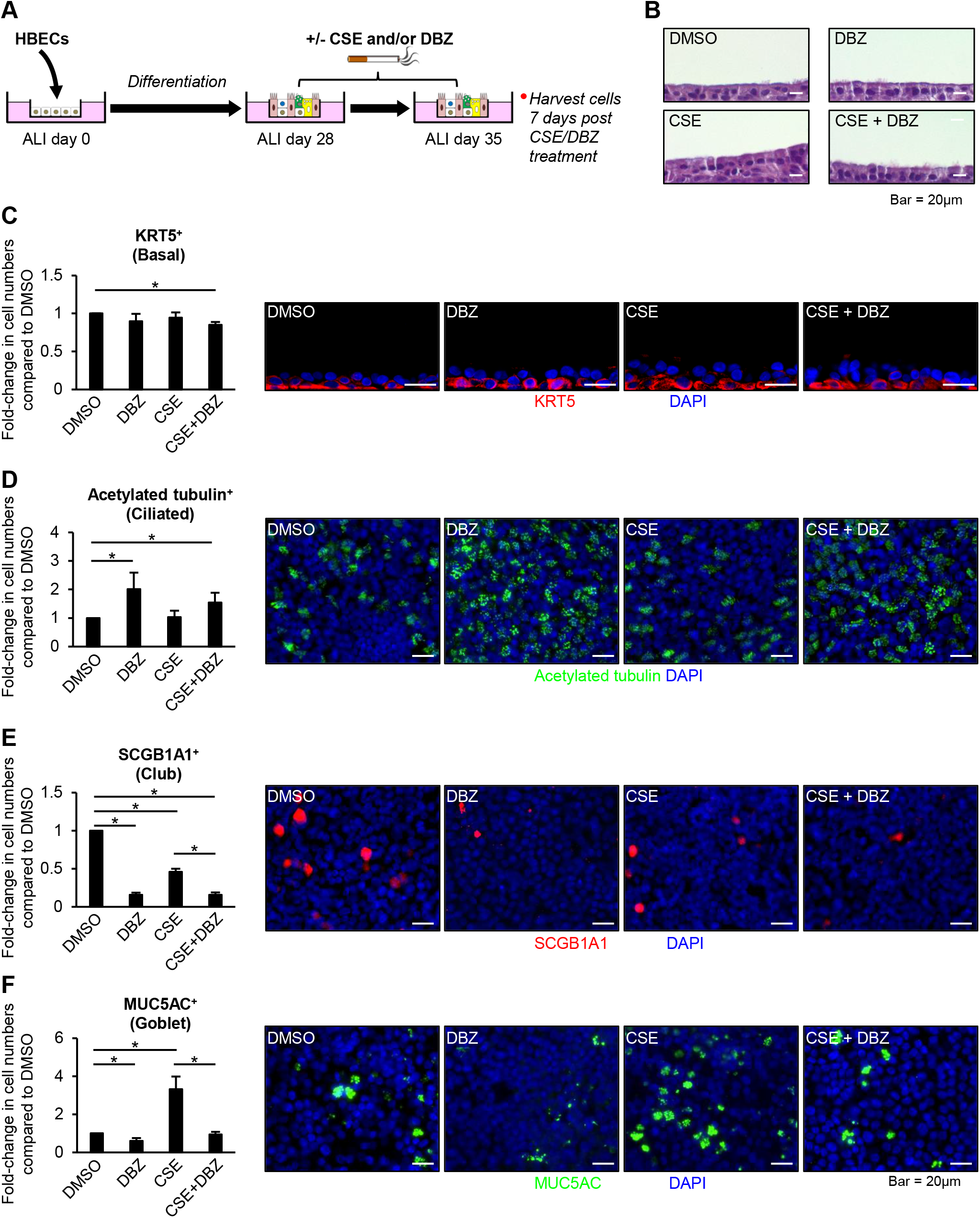
Treatment with the γ-secretase inhibitor Dibenzazepine (DBZ) suppresses CSE-induced goblet cell differentiation. **(A)** Schematic of air-liquid interface (ALI) model. Primary human bronchial epithelial cells (HBECs) from nonsmokers were cultured on ALI for 28 days to differentiate into a pseudostratified epithelium containing basal, ciliated and secretory (club and goblet) cells. At ALI day 28, the cultures were treated with DMSO (vehicle), DBZ (0.1 μM), DMSO + CSE (2.5%) and CSE + DBZ for 7 days. **(B)** H&E staining. **(C-F)** Immunofluorescent staining of basal cells (KRT5, red), ciliated cells (acetylated tubulin, green), club cells (SCGB1A1, red), and goblet cells (MUC5AC, green). Images were quantified using the ImageJ software (NIH). Data presented as mean fold-change in cell numbers compared to DMSO treated cells from n=4 independent experiments (see Figure E2 for presentation of these data as percentage positive cells for each condition). Scale bar = 20μm. Error bars indicate SEM. * p<0.05.

### CSE exposure promotes NOTCH3 signaling activation to regulate goblet cell differentiation

Previous studies have shown that signaling via the NOTCH1-3 receptors and the NOTCH ligands DLL1, JAG1 and JAG2 plays an important role in regulating differentiation of the adult airway epithelium in response to injury and environmental insult (12, 14, 15, 17–20, 22, 31–35). Therefore, we assessed the effect of CSE exposure on expression (mRNA levels) of these genes. No significant changes in expression were observed for DLL1, JAG1, NOTCH2 and NOTCH3 in response to CSE (Figure 4A). However, consistent with our RNA-Seq data (Figure 2B) a significant decrease in expression of JAG2 (0.64 fold) and NOTCH1 (0.56 fold) was observed in response to CSE (Figure 4A). Activation of canonical ligand-dependent NOTCH signaling results in proteolytic cleavage of the NOTCH receptor at the intracellular transmembrane region and subsequent release of the NOTCH intracellular domain (NICD) into the cytoplasm that can enter the nucleus to induce transcription (23). Therefore, to assess NOTCH signaling activation in response to CSE, we quantified the protein levels of the NICD from NOTCH1, 2 and 3 in whole cell lysates by western blot analysis. Consistent with the gene expression patterns of NOTCH1 and NOTCH2 (Figure 4A), CSE treatment resulted in a significant decrease in NOTCH1 NICD protein levels and no consistent change in NOTCH2 NICD protein levels compared to untreated cells (Figure 4B-C). In contrast to NOTCH1 and NOTCH2, a significant increase in NOTCH3 NICD protein levels was observed in CSE treated cells (Figure 4B-C), despite no change in NOTCH3 mRNA levels (Figure 4A). Confirming the western blot analysis, confocal immunofluorescent staining demonstrated increased NOTCH3 levels in both the cytoplasm and nucleus of CSE treated *vs.* untreated cells (Figure 4D). The increased nuclear staining of NOTCH3 NICD is indicative of increased NOTCH signaling, suggesting that CSE exposure might promote goblet cell differentiation via activation of NOTCH3-dependent signaling. To investigate the expression of NOTCH3 in goblet cells we performed co-localization of NOTCH3 with MUC5AC using immunofluorescence (Figure 4E). In both untreated and CSE treated cells, NOTCH3 was expressed in a subset of cells that surrounded areas of MUC5AC positive goblet cells. However, CSE treatment led to co-localization of NOTCH3 with some MUC5AC positive cells. To determine the ability of NOTCH3 signaling to initiate and promote goblet cell differentiation in the absence of CSE, primary HBECs from nonsmokers were infected with control lentivirus or lentivirus over-expressing NICD3 and harvested 7 days post infection (Figure 4E). Compared to control lentivirus infected cells, over-expression of NICD3 led to a significant increase in expression of SPDEF (2.47 fold) and MUC5AC (7.04 fold) demonstrating that NOTCH3 activation can promote goblet cell differentiation (Figure 4F). In addition, these data demonstrate that SPDEF is a downstream target of NOTCH3 signaling.

**Figure 4.**
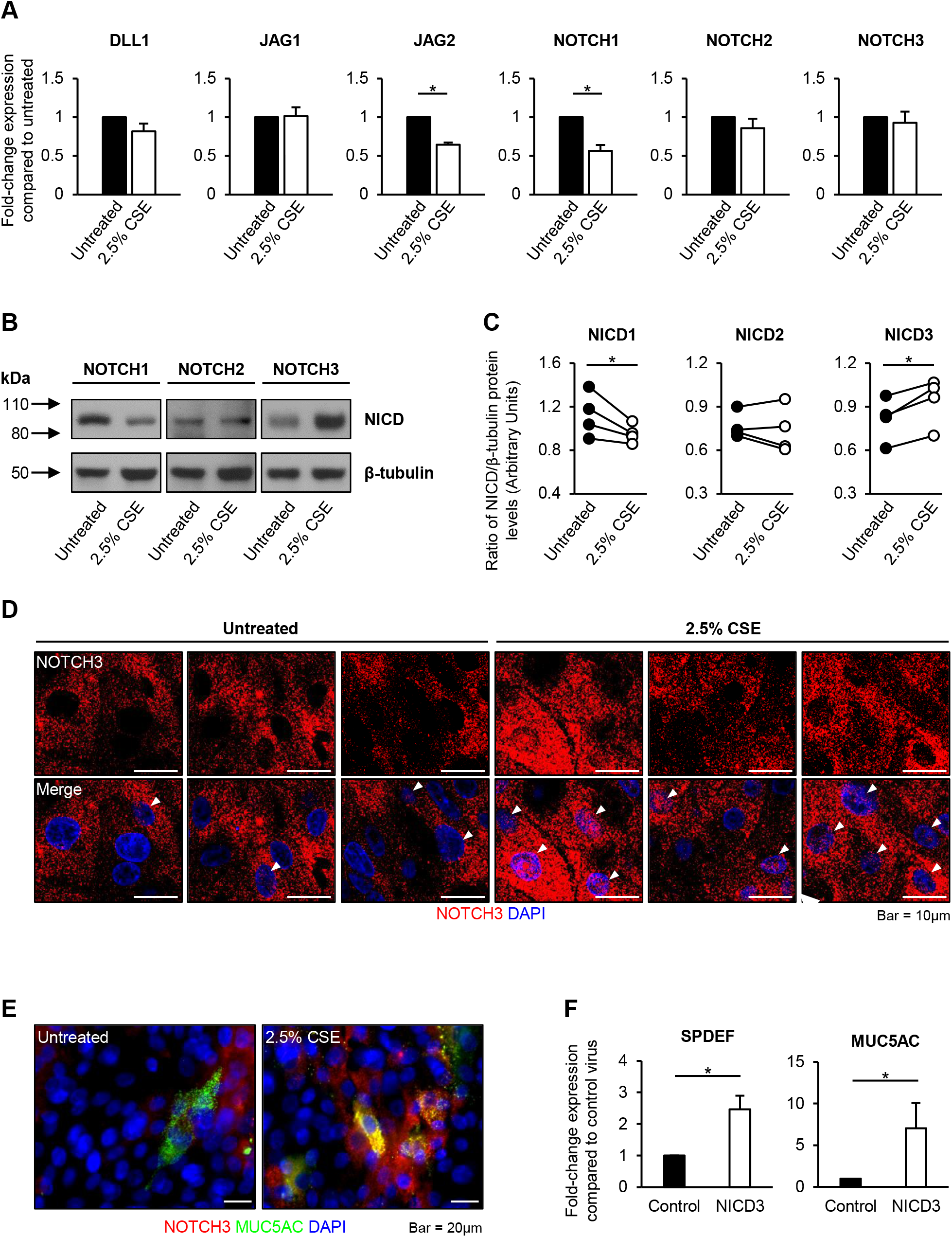
CSE treatment induces NOTCH3 activation. **(A-E)** Primary human bronchial epithelial cells (HBECs) from nonsmokers were cultured on ALI for 28 days to differentiate into a pseudostratified epithelium containing basal, ciliated and secretory (club and goblet) cells. At ALI day 28, the cultures were untreated or treated for 7 days with 2.5% cigarette smoke extract (CSE) then harvested for analysis. **(A)** qPCR of NOTCH ligand (DLL1, JAG1 and JAG2) and receptor (NOTCH1, NOTCH2 and NOTCH3) expression in untreated and CSE treated cells. Data presented as mean fold-change in expression compared to untreated cells from n=4 independent experiments. **(B)** Western analysis of the NOTCH1-3 receptor intracellular domain (NICD) in untreated and CSE treated cells. β-tubulin was used a loading control. **(C)** Quantification of NOTCH1-3 NICD protein levels. Data presented as ratio of NICD/β-tubulin protein levels between untreated and CSE treated cells from n=4 independent experiments. **(D)** Immunofluorescent staining and confocal microscopy of NOTCH3 (red) and nuclei (blue, DAPI) in untreated and CSE treated cells. Scale bar = 10μm. White arrows indicate nuclear staining of NOTCH3. **(E)** Immunofluorescent staining of NOTCH3 (red), MUC5AC (green) and nuclei (blue, DAPI) in untreated and CSE treated cells. Scale bar = 20μm. **(F)** Primary HBECs from nonsmokers were infected on ALI with either control lentivirus (Lenti-Control) or lentivirus expressing the activated NOTCH3 NICD (Lenti-NICD3). Following 7 days of culture (ALI day 7) the cells were harvested. qPCR of SPDEF and MUC5AC expression in Lenti-Control and Lenti-NICD3 infected cells. Data presented as mean fold-change in expression compared to Lenti-Control cells in n=6 independent experiments (single ALI well analyzed for gene expression per experiment). For panels **A**, **C** and **F** the error bars indicate SEM. * p<0.05, ** p<0.005.

### siRNA-mediated knockdown of NOTCH3 suppresses CSE-mediated induction of goblet cell differentiation

Since CSE induces NOTCH3/NICD3 activation (Figure 4B-D) and over-expression of NICD3 induces goblet cell differentiation (Figure 4F), we next tested the role of NOTCH3 signaling in regulating CSE-dependent induction of goblet cell differentiation. Due to difficulties in achieving high knockdown efficiency with siRNA in fully differentiated airway epithelium (e.g. ALI day 28) we adopted an alternative approach. Primary HBECs from nonsmokers were transfected with either control siRNA or NOTCH3 specific siRNA at the time of seeding the cells on ALI culture and then treated with CSE for 7 days from ALI day 0 to 7. Quantitative PCR analysis was then performed to measure NOTCH3 knockdown efficiency, response to CSE (CYP1A1 expression) and goblet cell differentiation (SPDEF and MUC5AC expression). Due to the variability in differentiation kinetics and response to CSE between the different HBEC donors at early time points on ALI, the data are presented as individual experiments (Figure 5). Compared to siRNA Control transfected cells, we consistently observed >70% knockdown of NOTCH3 expression in siRNA NOTCH3 transfected cells in the absence and presence of CSE. Across three independent experiments, CSE treatment of siRNA Control transfected cells resulted in increased expression of the oxidative-response gene CYP1A1 suggesting that the cells were responding to CSE (Figure 5). Furthermore, CSE exposure induced goblet cell differentiation as assessed by increased expression of SPDEF and MUC5AC. Similar to siRNA Control cells, CSE treatment of siRNA NOTCH3 transfected cells increased CYP1A1 expression to comparable levels. However, knockdown of NOTCH3 reduced CSE-induced goblet cell differentiation as evidenced by decreased expression of SPDEF and MUC5AC. Combined, these data suggest that CSE exposure induces goblet cell differentiation in the human airway epithelium via activation of NOTCH3 signaling.

**Figure 5.**
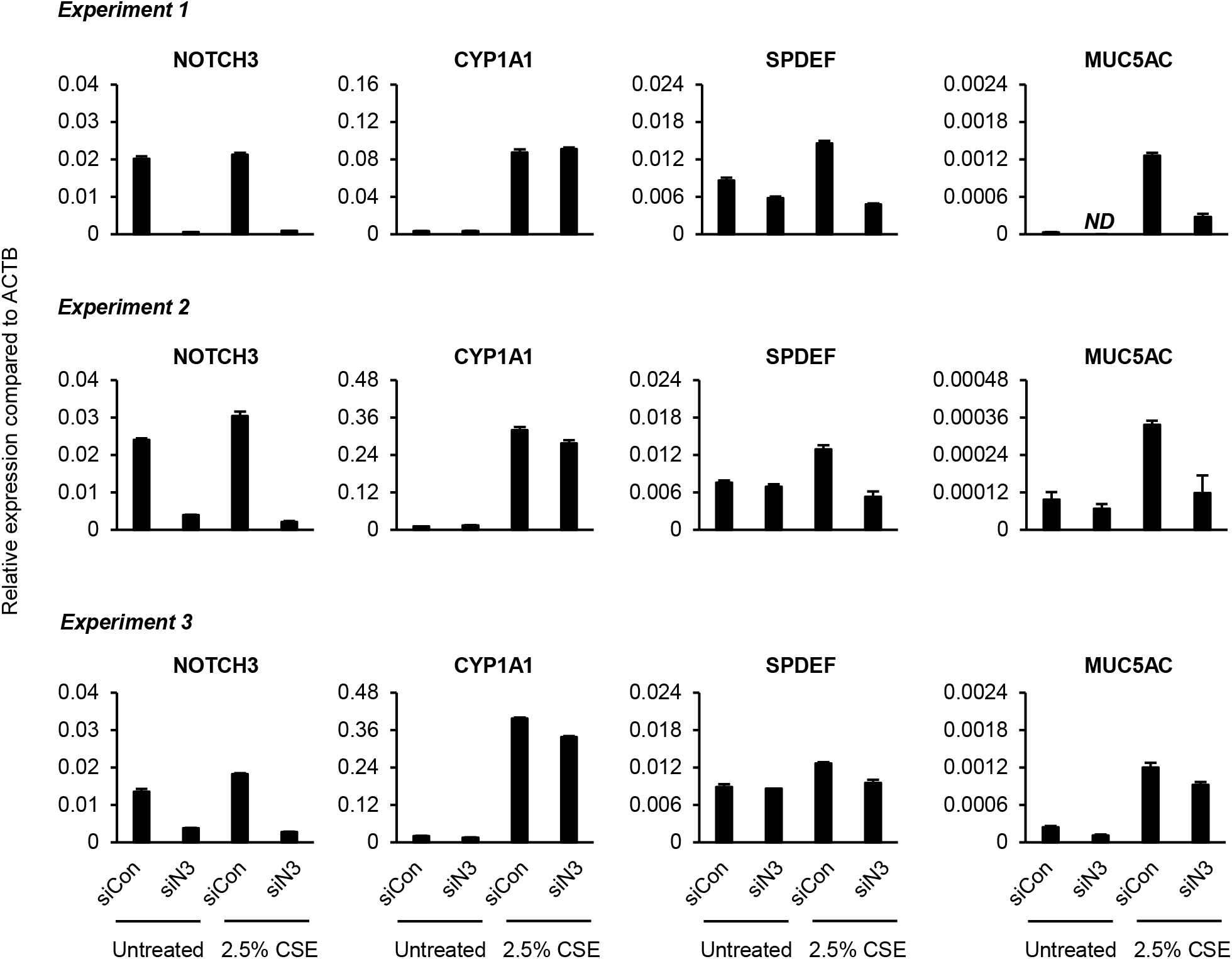
siRNA-mediated knockdown of NOTCH3 suppresses CSE-dependent induction of SPDEF and MUC5AC. **(A-C)** Primary human bronchial epithelial cells (HBECs) from nonsmokers (n=3 independent donors) were either transfected with control (siCon) or NOTCH3 (siN3) specific siRNA during seeding on ALI culture. At ALI day 0 the cultures were either untreated or treated for 7 days with 2.5% cigarette smoke extract (CSE) then harvested for qPCR analysis of NOTCH3, CYP1A1, SPDEF and MUC5AC. For each donor (Experiment 1-3) the data is presented as mean relative expression compared to ACTB from n=3 ALI wells. Error bars indicate SEM. ND indicates “not detected”.

### CSE exposure activates NOTCH3 signaling in COPD cells to regulate goblet cell differentiation

Alterations in NOTCH pathway components have been identified in the airways of smokers with and without COPD relative to healthy nonsmokers at the mRNA, protein, and epigenetic levels (13, 32, 36, 37). Furthermore, HBECs from COPD patients have been shown to have an altered differentiation capacity compared to healthy controls (38, 39). Therefore, from a therapeutic perspective it is critical to determine if CSE exposure impacts goblet cell differentiation in COPD airway epithelium in a similar manner to healthy nonsmokers. To this end, HBECs from COPD smokers were cultured *in vitro* on ALI for 28 days and then exposed to 2.5% CSE for 7 days (Figure 6). Whilst all the COPD HBEC donors differentiated into a pseudostratified epithelium, comparison with nonsmoker HBEC cultures at ALI day 35 demonstrated the COPD airway epithelium had lower numbers of ciliated, club and goblet cells (Figure E3). However, CSE exposure of COPD airway epithelium led to the appearance of a remodeled epithelium (Figure 6A) and a significant increase in expression of the oxidative-stress response gene CYP1A1 (102.5 fold) (Figure 6B) similar to that observed in nonsmoker airway epithelium (Figure 2B-C). Expression of the basal (KRT5) and ciliated (DNAI1) cell markers did not change in the presence of CSE, whereas expression of the club cell marker SCGB1A1 was significantly decreased (0.29 fold) with a corresponding significant increase in expression of the goblet cell markers MUC5AC (17.8 fold) and SPDEF (4.1 fold) (Figure 6B). The CSE-dependent changes in cell-type specific marker gene expression were then validated by quantifying the number of basal, ciliated, club and goblet cells histologically (Figure 6C). Corroborating the gene expression data, there were no significant changes in the numbers of basal (KRT5^+^) and ciliated (acetylated tubulin^+^) cells in CSE treated ALI cultures compared to untreated cultures. Furthermore, a significant decrease in the numbers of SCGB1A1^+^ club cells (0.38 fold) and a corresponding significant increase in MUC5AC^+^ goblet cells (2.97 fold) was observed with CSE treatment (Figure 6C). To characterize the CSE-dependent transcriptome changes in COPD airway epithelium, RNA-Seq was performed on untreated or 2.5% CSE treated cells 7 days post treatment (ALI day 35). Our analysis identified 230 genes (140 up-regulated and 90 down-regulated) with significant (FDR adjusted p<0.05) expression changes (File E3). Molecular pathways associated with the 230 CSE-responsive genes included those previously associated with cigarette smoking and COPD, including: LXR/RXR activation (14 genes), IL-17A signaling in airway cells (8 genes), aryl hydrocarbon receptor signaling (10 genes), nicotine degradation II (6 genes) and xenobiotic metabolism signaling (11 genes) (File E4) (24, 25). Comparison of the 230 CSE-responsive genes identified in COPD airway epithelium with the 273 CSE-responsive genes identified in nonsmoker airway epithelium revealed 125 genes common to both groups (File E5). Within this 125 gene set were many of the genes associated with goblet cell biology altered in nonsmoker cells (Figure 2B) including inflammatory mediators (CXCL3, CXCL5, IL1β, IL1RN and IL19), endoplasmic reticulum stress (CREB3L1 and ERN2), mucus hypersecretion (SLC12A2 and SLC26A4) and NOTCH signaling (NOTCH1 and SPDEF) (Figure E5). Importantly, the direction of CSE-induced gene expression changes for these genes was conserved in nonsmoker and COPD airway epithelium (Figure 2B and Figure E5) suggesting a shared mechanism of CSE-dependent goblet cell differentiation.

**Figure 6.**
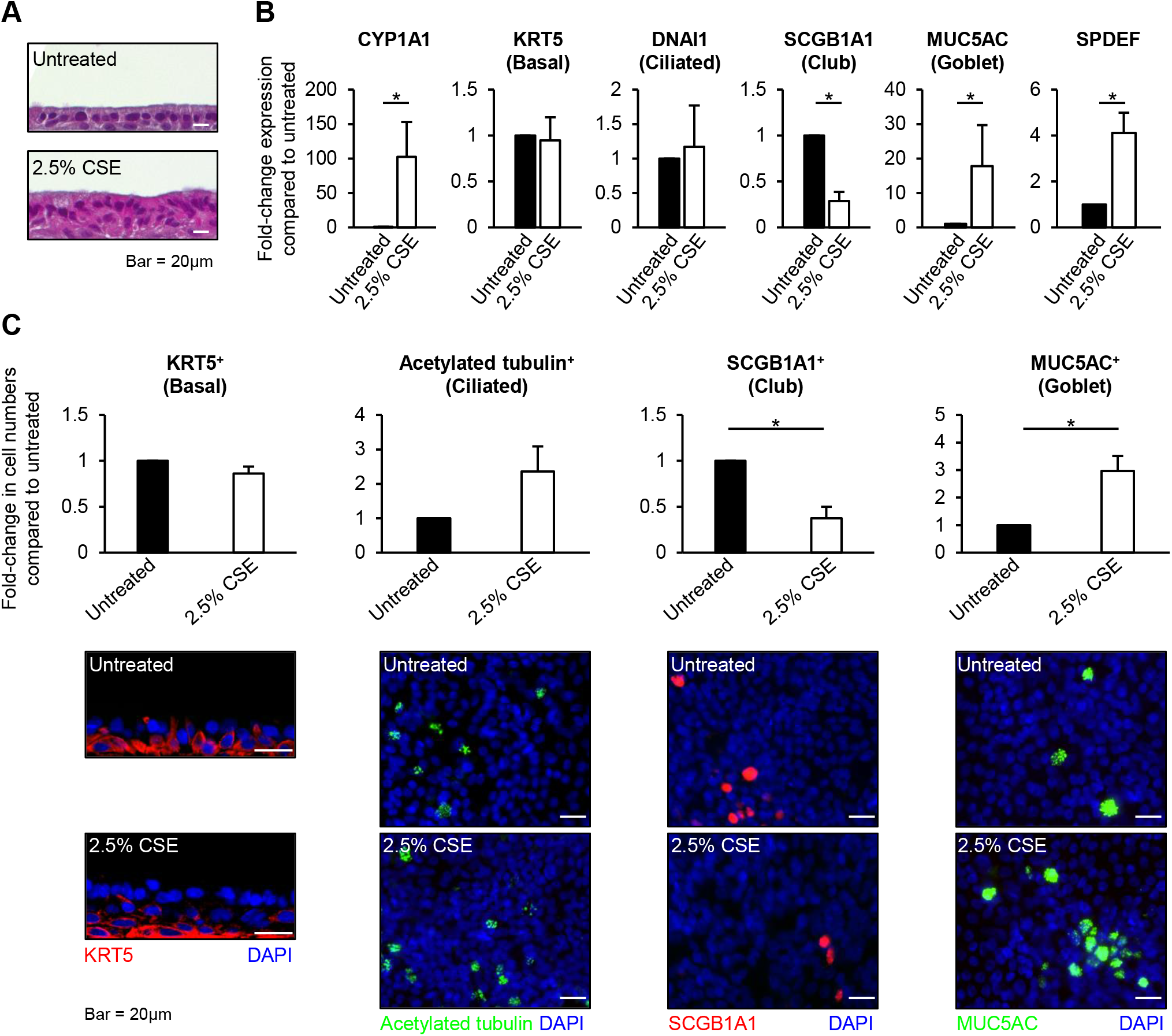
*In vitro* CSE exposure of COPD airway epithelium induces goblet cell differentiation. **(A-C)** Primary human bronchial epithelial cells (HBECs) from COPD smokers were cultured on ALI for 28 days to differentiate into a pseudostratified epithelium containing basal, ciliated and secretory (club and goblet) cells. At ALI day 28, the cultures are then untreated or treated for 7 days with 2.5% cigarette smoke extract (CSE) before harvest. **(A)** H&E staining. Scale bar = 20μm. **(B)** qPCR of CYP1A1, KRT5, DNAI1, SCGB1A1, MUC5AC and SPDEF expression in untreated and CSE treated cells. Data presented as mean fold-change in expression compared to untreated cells from n=4 independent experiments. **(C)** Immunofluorescent staining of basal cells (KRT5, red), ciliated cells (acetylated tubulin, green), club cells (SCGB1A1, red), and goblet cells (MUC5AC, green). Images were quantified using the ImageJ software (NIH). Data represented as mean fold-change in cell numbers compared to untreated cells from n=4 independent experiments (see Figure E4 for presentation of these data as percentage positive cells for each condition). Scale bar = 20μm. For panels **B** and **C** the error bars indicate SEM. * p<0.05.

Based on our demonstration that CSE induces goblet cell differentiation in nonsmoker cells via a NOTCH-dependent manner, we quantified gene expression of the NOTCH ligands (DLL1, JAG1 and JAG2) and receptors (NOTCH1, NOTCH2 and NOTCH3) in CSE treated *vs.* untreated COPD airway epithelium. Similar to nonsmoker airway epithelium (Figure 4A), no significant changes in expression were observed for DLL1, JAG1, NOTCH2 and NOTCH3 (Figure 7A), whereas CSE significantly decreased expression of JAG2 (0.63 fold) and NOTCH1 (0.66 fold) (Figure 7A). We next investigated NICD protein levels from NOTCH1, 2 and 3 in response to CSE by western blot analysis (Figure 7B-C). Consistent with the results from nonsmoker airway epithelium (Figure 4B-C), CSE treatment lead to increased protein levels of NICD3 and decreased levels of NOTCH1 NICD (Figure 7B-C). However, we also observed a decrease in NOTCH2 NICD which was also observed in a subset of nonsmoker donors (Figure 4C).To determine the role of NOTCH3 signaling in regulating CSE-dependent induction of goblet cell differentiation in COPD airway epithelial cells, we performed siRNA mediated knockdown of NOTCH3 on ALI culture in the absence and presence of CSE using the same strategy described for nonsmoker HBECs. Due to variability in differentiation kinetics and response to CSE between the different COPD HBEC donors, the data are presented as individual experiments. Compared to siRNA Control transfected cells, we consistently observed >90% knockdown efficiency of NOTCH3 expression in siRNA NOTCH3 transfected cells in the absence and presence of CSE. From two independent experiments, CSE treatment of cells transfected with either siRNA Control or siRNA NOTCH3 resulted in increased expression of the oxidative-response gene CYP1A1 (Figure 7D). We were unable to reliably detect MUC5AC in untreated cells at ALI day 7, however, CSE exposure induced goblet cell differentiation with increased expression of SPDEF and MUC5AC (Figure 7D). Moreover, knockdown of NOTCH3 suppressed CSE-dependent induction of the goblet cell associated genes SPDEF and MUC5AC (Figure 7D). Combined, these data suggest that CSE exposure of COPD airway epithelium induces goblet cell differentiation in a NOTCH3-dependent manner.

**Figure 7.**
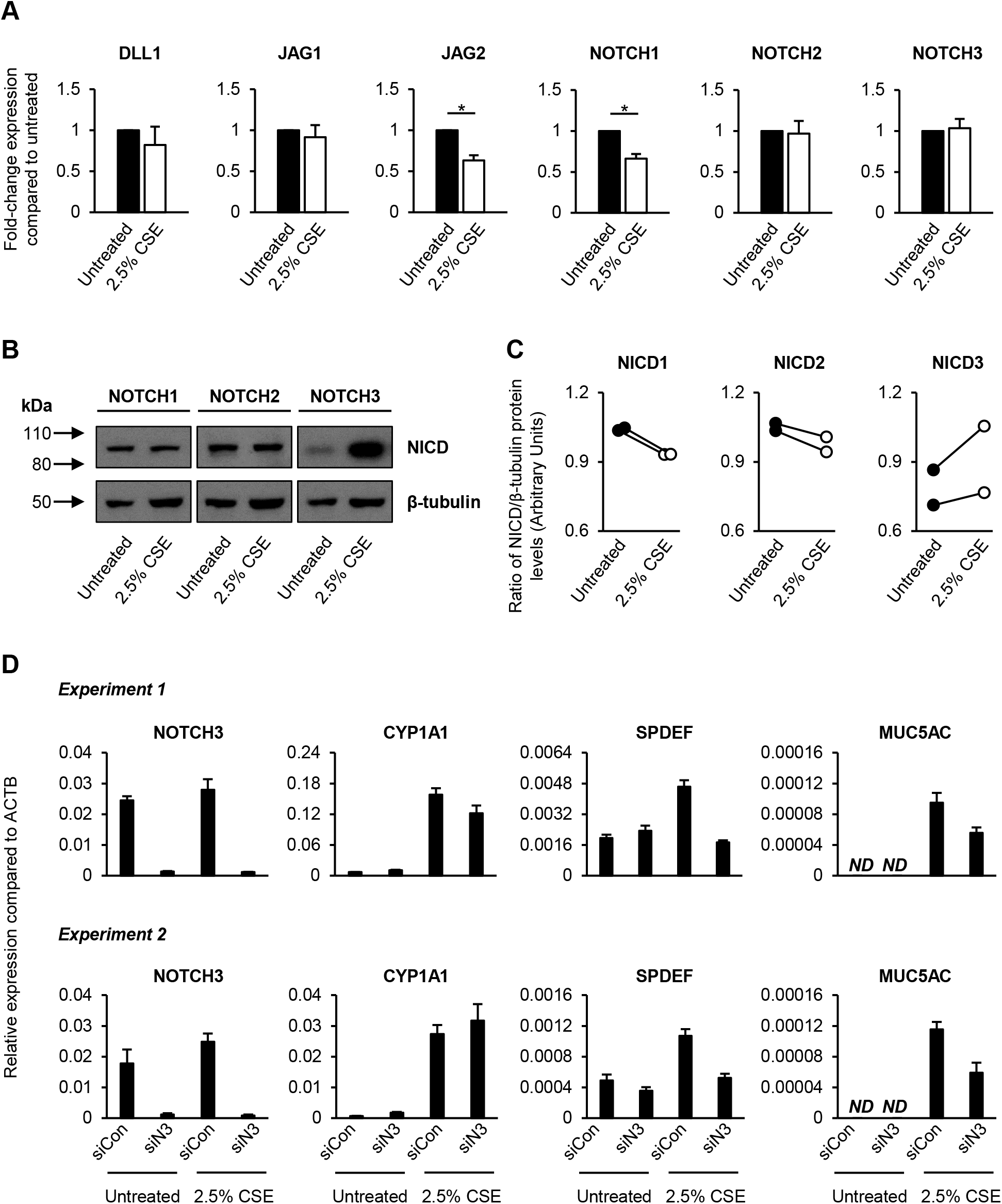
CSE induces SPDEF and MUC5AC expression in COPD airway epithelium in a NOTCH3-dependent manner. **(A-C)** Primary human bronchial epithelial cells (HBECs) from COPD smokers were cultured on ALI for 28 days to differentiate into a pseudostratified epithelium containing basal, ciliated and secretory (club and goblet) cells. At ALI day 28, the cultures were untreated or treated for 7 days with 2.5% cigarette smoke extract (CSE) then harvested for analysis. **(A)** qPCR of NOTCH ligands (DLL1, JAG1 and JAG2) and receptors (NOTCH1, NOTCH2 and NOTCH3) expression in untreated and CSE treated cells. Data presented as mean fold-change in expression compared to untreated cells from n=4 independent experiments. Error bars indicate SEM. * p<0.05. **(B)** Western analysis of the NOTCH1-3 receptor intracellular domain (NICD) in untreated and CSE treated cells. β-tubulin was used a loading control. **(C)** Quantification of NOTCH1-3 NICD protein levels. Data presented as ratio of NICD/β-tubulin protein levels between untreated and CSE treated cells from the experiment shown in panel **B**. **(D)** Primary HBECs from COPD smokers (n=2 independent donors) were either transfected with control (siCon) or NOTCH3 (siN3) specific siRNA during seeding on ALI culture. At ALI day 0 the cultures were either untreated or treated for 7 days with 2.5% cigarette smoke extract (CSE) then harvested for qPCR analysis of NOTCH3, CYP1A1, SPDEF and MUC5AC. For each donor (Experiment 1-2) the data is presented as mean relative expression compared to ACTB from n=3 ALI wells. Error bars indicate SEM. ND indicates “not detected”.

## Discussion

The NOTCH signaling pathway plays a crucial role in regulating cell fate decisions of multiple cell types in the human and murine airway epithelium including basal, ciliated, club and goblet cells (11). Here we demonstrate that CSE exposure of airway epithelium generated from nonsmoker and COPD-smoker derived HBECs on ALI culture promotes GCMH in a NOTCH-dependent manner. Moreover, treatment with a NOTCH pathway/γ-secretase inhibitor (DBZ) suppressed CSE-dependent induction of goblet cell differentiation indicating that CSE activates NOTCH signaling to induce the GCMH phenotype. Further analysis of specific NOTCH components revealed that CSE exposure increased NOTCH3 receptor intracellular domain (NICD3) protein levels and nuclear localization indicative of increased NOTCH3 activation. Moreover, constitutive over-expression of NICD3 was sufficient to induce SPDEF and MUC5AC expression even in the absence of CSE, whereas siRNA mediated knockdown of NOTCH3 in differentiating nonsmoker and COPD smoker HBECs suppressed CSE-mediated induction of SPDEF and MUC5AC. Overall, these data demonstrate that activation of NOTCH3 signaling regulates airway epithelial goblet cell differentiation in response to CSE, leading to enhanced goblet cell differentiation and development of GCMH.

It is well established that the NOTCH pathway plays a critical role in regulating goblet cell differentiation of the airway epithelium in both humans and mice during homeostasis and in response to injury or environmental insult (12–22). Previous studies have shown that interactions between JAG1-2 ligands and NOTCH1-3 receptors regulate development of GCMH in the asthmatic airway epithelium and in response to inflammatory cytokines, allergens, and viral infections (12, 15, 17–19), while their role in the CSE-induced GCMH model was unknown. Reid et al., (12) recently showed that NOTCH3 expression is upregulated in the airway epithelium of asthmatics and regulates MUC5AC expression, whereas Jing et al., (17) demonstrated that rhinovirus infection regulates development of GCMH in COPD airway epithelial cells in a NOTCH3-dependent manner. Our study expands on these findings by demonstrating that CSE exposure activates NOTCH3 signaling to induce goblet cell differentiation and MUC5AC expression in both nonsmoker and COPD airway epithelial cells.

Basal cells are the stem/progenitor cells of the airway epithelium in both humans and mice, and are capable of generating club cells, which in turn can undergo differentiation into goblet or ciliated cell lineages (5, 6). The NOTCH-dependent differentiation of basal cells into a pseudostratified airway epithelium is a multi-step process, with cell fate decisions involving a coordinated response between different combinations of NOTCH ligand-receptor signaling events (11, 14, 15, 20, 22, 32, 35, 40). Recent single cell RNA-sequencing studies have shed light on the cell type specific expression pattern of NOTCH signaling molecules (41) and shown that NOTCH3 expression is predominantly enriched within a sub-population of basal cells (termed supra-basal cells) and club cells. Work by Mori et al. (32) demonstrated that NOTCH3 activation leads to priming of basal cell differentiation into early/intermediate club cells, which in turn can differentiate into goblet cells or ciliated phenotypes through secondary NOTCH signaling. Previous work from our lab demonstrated that constitutive activation of NOTCH3 signaling *via* NICD3 increases both club and goblet cell differentiation of nonsmoker HBECs on ALI culture (14). In addition, work from Reid et al (12) showed that NOTCH3 activation regulates MUC5AC expression. Our data demonstrate that CSE exposure of ALI day 28 differentiated airway epithelium leads to increased numbers of goblet cells (MUC5AC^+^) and decreased numbers of club cells (SCGB1A1^+^), suggesting that CSE-dependent activation of NOTCH3 signaling in club cells may promote further differentiation of this cell population into goblet cells. Furthermore, siRNA mediated knockdown of NOTCH3 prevented CSE mediated induction of goblet cell differentiation. Further studies are ongoing to clarify how CSE impacts the differentiation of individual cell populations and the NOTCH3 signaling events that regulate this process.

Prior studies identified alterations in NOTCH signaling components at the mRNA, protein, and epigenetic levels in the small airway epithelium of smokers with and without COPD relative to nonsmokers (13, 32, 36, 37). A study by Tilley et al. (36) reported decreased NOTCH3 mRNA in the small airway epithelium in smokers *vs.* nonsmokers suggesting that NOTCH signaling was suppressed in response to smoking. However, changes in the protein levels of active NOTCH3 receptor (NICD3) were not assessed. In support of our study, cigarette smoke increased the levels of NOTCH3 protein in human lung adenocarcinoma both *in vitro* and *in vivo* (42). Furthermore, 24 weeks of cigarette smoke exposure of mice increased NOTCH signaling activity in lung lymphoid tissue (43). Importantly, the mechanism by which CSE activates NOTCH3 signaling to induce goblet cell differentiation remains unknown. Canonical NOTCH signaling is dependent on cell-cell contact and requires binding of a ligand on one cell to the receptor on a neighboring cell (23). Studies in both mice and humans identified the JAG1 and JAG2 ligands as the major regulators of cell differentiation in airway epithelium (18, 19, 22, 32, 34, 40). However, in our study CSE decreased expression of JAG2 in airway epithelium, but had no effect on JAG1 expression. Prior work from our lab showed that over-expression of JAG1 in an immortalized airway basal epithelial cell line led to increased secretory cell differentiation (22). Therefore, one possibility is that decreased JAG2 expression in response to CSE exposure leads to increased availability of JAG1 and subsequent activation of JAG1-NOTCH3-dependent signaling to promote goblet cell differentiation.

Cigarette smoke exposure may also regulate NOTCH3 activation via non-canonical ligand-independent mechanisms (23). It is noteworthy that CSE exposure increased NICD3 protein levels (and nuclear localization) with no change in expression of NOTCH3 mRNA, suggesting that CSE regulates NOTCH3 protein levels post-transcriptionally. Direct interaction of post-secretase cleaved NOTCH3 fragments with the E3-ubiquitin ligase WWP2 leads to mono-ubiquitination of NICD3, which promotes its sorting to and degradation in lysosomes leading to suppression of NOTCH3 signaling (44). Therefore, CSE exposure may induce NOTCH3 activation via increasing NICD3 half-life by preventing its ubiquitination and subsequent degradation in lysosomes. While we do not observe any changes in WWP2 expression in response to CSE, it is plausible that CSE alters the expression or activity of other E3-ubiquitin ligases that regulate NICD3 ubiquitination and stability. Alternatively, cellular redox status has been shown to regulate NOTCH3 protein levels through lysosome-dependent protein degradation (45). Treatment of cancer cells with the antioxidant N-acetyl-cysteine leads to increased NOTCH3 degradation through lysosome-dependent pathways with no change in NOTCH3 mRNA levels (46). Moreover, cigarette smoke exposure induces high levels of oxidative stress due to the cumulative effect of intrinsic reactive oxygen species (ROS) present in the smoke, and increased cellular ROS production (47, 48). Based on this knowledge, further studies are warranted to evaluate if cigarette smoke induced oxidative stress in the airway epithelium leads to impairment of lysosome-dependent NOTCH3 protein degradation.

Despite the knowledge that NOTCH3 signaling plays an important role in regulating airway epithelial differentiation (12, 14, 17, 32), the downstream signaling events and specific targets are largely unknown. Our demonstration that over-expression of NICD3 leads to increased expression of the goblet cell promoting transcription factor SPDEF (30) in HBECs, suggests that NOTCH3 regulates goblet cell differentiation at least in part via SPDEF-dependent mechanisms. Our RNA-Seq analysis in both nonsmoker and COPD smoker airway epithelial cells showed that CSE exposure also increased the expression of other genes associated with goblet cell biology and chronic lung disease including growth factors (e.g. AREG), genes associated with endoplasmic reticulum stress (e.g. ERN2) and inflammation (e.g. IL1β). Interleukin-1β (IL1β) is a pro-inflammatory cytokine up-regulated in the small airway epithelium of COPD patients (49) and in the bronchoalveolar lavage fluid isolated from cigarette smoke exposed mice (48). Additionally, *in vitro* stimulation of HBECs or administration of mice with recombinant IL1β induces goblet cell differentiation with increased SPDEF, ERN2 and MUC5AC expression (26, 50). Therefore, in addition to the direct induction of SPDEF by NOTCH3 activation, upregulation of IL1β expression in response to CSE may further contribute to goblet cell differentiation in a positive feedback cycle by induction of SPDEF expression in either an autocrine or paracrine manner.

In summary, our data demonstrate that CSE induced NOTCH3 activation is a key regulator of GCMH development in the airway epithelium derived from both nonsmokers and COPD subjects. Thus, targeting NOTCH3 signaling could be used as a novel therapeutic strategy to control GCMH in smokers with and without COPD.

## Acknowledgements

The authors would like to thank Drs. Linda Thompson, Dean Dawson, Lorin Olson and Xiao-Hong Sun at the Oklahoma Medical Research Foundation (OMRF) for discussions, guidance, and support. We also thank the Imaging Core Facility at OMRF for assistance in sample processing and confocal microscopy.

## Abbreviations

ALI: Air liquid interphase
AQP5: Aquaporin 5
AREG: Amphiregulin
ATP12A: ATPase, H+/K+ transporting, nongastric, alpha polypeptide
CLCA2: Chloride channel accessory 2
COPD: Chronic obstructive pulmonary disease
CREB3L1: CAMP responsive element binding protein 3 like 1
CSE: Cigarette smoke extract
CXCL3/5: C-X-C motif chemokine ligand 3/5
CYP1A1: Cytochrome P450, family 1, subfamily A, polypeptide 1
DBZ: Dibenzazepine
DLL1/3/4: Delta-like protein 1/3/4
ERN2: Endoplasmic reticulum to nucleus signaling 2
FDR: False discovery rate
GCMH: Goblet cell metaplasia and hyperplasia
HBEC: Human bronchial epithelial cell
IL1RN: Interleukin 1 receptor antagonist
IL1β: Interleukin 1β
JAG 1/2: Jagged-1/2
NICD: Notch intracellular domain
ROS: Reactive oxygen species
SLC12A2: Solute carrier family 12 member 2
SLC26A4: Solute carrier family 26 member 4
SLC31A1: Solute carrier family 31 member 1
SPDEF: Sam pointed domain-containing Ets transcription factor
WWP2: WW domain-containing protein 2

## Online Data Supplement

### Extended Materials and Methods

#### Primary human bronchial epithelial cell (HBEC) culture

Primary human bronchial epithelial cells (HBECs) were purchased (Lonza, Morristown, NJ, USA) from nonsmokers (Cat# CC-2540) and COPD smokers (Cat# 00195275 and 00195275S). The demographic details of all the donors is summarized in Supplementary Table I. The cells were maintained in BronchiaLife™ epithelial airway medium (BLEAM) (Cat# LL-0023, Lifeline^®^ Cell Technology, Oceanside, CA, USA) supplemented with Penicillin (100 I.U./ml) - Streptomycin (100 μg/ml) under standard cell culture conditions as previously described (E1). All experiments were performed with either Passage 2 or 3 cells.

#### Air-liquid interface (ALI) culture

Primary HBECs were differentiated using air-liquid interface (ALI) culture to generate a pseudostratified epithelium consisting of basal, ciliated and secretory (club and goblet) cells, using a previously described protocol (E1) with the following modifications. Briefly, 1 × 10^5^ cells in 100 μl of BLEAM media were seeded in the apical chamber of a 0.4 μm pore-sized Transwell® insert (Cat# 3470, Corning^®^, Corning, NY, USA) pre-coated with human type IV collagen (Cat# C7521, Sigma Aldrich, St. Louis, MO, USA) with 1 ml of BLEAM in the basolateral chamber (ALI day - 2). The following day, fresh BLEAM media was replaced in the apical and basolateral chambers (100 μl and 1 ml, respectively). Following two days of submerged culture, media from the apical chamber was removed to expose the cells to air (ALI day 0), and 1 ml of HBTEC ALI differentiation medium (Cat# LM-0050, Lifeline^®^ Cell Technology) supplemented with Penicillin (100 I.U./ml) - Streptomycin (100 μg/ml) was added to the basolateral chamber. The media was replaced every other day and the cells allowed to differentiate for up to 28 days.

#### Preparation of cigarette smoke extract (CSE)

The smoke generated from two 3R4F Reference cigarettes (Center for Tobacco Reference Products, University of Kentucky College of Agriculture, Lexington, KY, USA) was bubbled through 30 ml of HBTEC ALI media (without antibiotics) in a glass apparatus (Cat# 1760-250, PYREX^®^, Oneonta, NY, USA), at the rate of 1 puff/minute, with each puff lasting 2-3 seconds. The extract was allowed to sit for 30 minutes in the glass apparatus, and then filtered through a 0.2 μm filter. The consistency of the CSE was standardized by measuring the OD of the CSE at 320 nm. An OD of 2.2 was obtained after burning two cigarettes and was considered 100% CSE. The CSE was checked to confirm a pH of 7.4 and then aliquoted into 1 ml single-use aliquots and stored in −80°C until further use. Further dilutions were made in complete HBTEC ALI media.

#### CSE and DBZ treatments

Cells cultured on ALI were treated with non-toxic concentrations of CSE (up to 2.5%) via the basolateral chamber for 7 days (either ALI day 0-7 or ALI day 28-35) with freshly thawed CSE added at each media change. For experiments involving suppression of NOTCH signaling, the cells were treated with CSE and/or DBZ (0.1 μM, Cat# 565789, EMD Chemicals Inc., Gibbstown, NJ, USA) from ALI day 28 to 35 with an equal volume of DMSO (Sigma Aldrich) used as vehicle control.

#### RNA extractions and Real-time qPCR analysis

Total RNA was extracted via direct lysis of cells in the ALI well using the PureLink™ RNA mini kit (Cat# 12183018A, Thermo Scientific, Waltham, MA, USA) and included DNase treatment (Cat# 12185-010, Thermo Scientific) on the column to remove contaminating genomic DNA. RNA concentrations were measured with a NanoPhotometer® N60 (Implen, Westlake Village, CA, USA) and cDNA was generated from 250 ng of total RNA per sample using random hexamers (Applied Biosystems™ High Capacity cDNA Reverse Transcription Kit, Cat# 4374966, Thermo Scientific) and subsequent quantitative PCR (qPCR) analysis performed using iTaq™ Universal SYBR® Green supermix (Cat# 1725124, Bio-Rad, Hercules, CA, USA) on the Bio-Rad CFX96 Touch™ Real-Time PCR system. All samples were analyzed in duplicate with relative expression levels determined using the dCt method with ACTB (β-actin) as the endogenous control. The following gene-specific primers were purchased from Bio-Rad (PrimePCR™, Bio-Rad): ACTB (qHsaCED0036269), CYP1A1 (qHsaCID0010608), KRT5 (qHsaCED0047798), DNAI (qHsaCID0017936), SCGB1A1 (qHsaCID0018013), MUC5AC (qHsaCID0017663), SPDEF (qHsaCID0021209), DLL1 (qHsaCID0011257), JAG1 (qHsaCID0006831), JAG2 (qHsaCED0047702), NOTCH1 (qHsaCID0011825), NOTCH2 (qHsaCED0005739) and NOTCH3 (qHsaCID0006529), and the assays performed using the manufacturer’s recommend cycling parameters. Unless indicated, gene expression levels were assessed in 3 replicate ALI wells for each time point and condition within every independent experiment. The mean of these 3 wells was then calculated as the final expression value.

#### Histological and Immunofluorescence staining

Cells were fixed directly in the ALI wells with 10% neutral buffered formalin (NBF) for 20 minutes at room temperature (RT), washed with phosphate buffered saline (PBS) and stored at 4°C until further use. Due to the reversible nature of NBF fixation, the wells were re-fixed if not used within 3 weeks. The fixed cells were either paraffin embedded and cross-sectioned into 5 μm sections (ALI sections) or used directly for top-staining. Images of hematoxylin and eosin (H&E) stained cross-sections were taken using an Olympus BX40 brightfield microscope (Olympus Corporation, Shinjuku City, Tokyo, Japan). Immunofluorescent (IF) staining of the paraffin embedded sections was performed using the following protocol. First, the sections were baked in a dry oven at 65°C for 30 minutes and allowed to cool down for another 15 minutes. Next, the sections were deparaffinized by dipping the slides in different solutions in the following order: two chambers of Xylene; two chambers of 100% Ethanol; two chambers of 95% Ethanol; one chamber of 70% Ethanol; one chamber of deionized water. The sections were then subjected to heat-induced antigen retrieval by dipping in 1X sodium citrate buffer (Cat# AP-9003-125, Thermo Scientific), and placed in a commercially available antigen retrieval system (2100 Antigen Retriever, BioVendor LLC, Asheville, NC, USA). After completion of the antigen retrieval cycle, the slides were washed briefly in running tap water, followed by blocking with 10% goat serum in PBS, for 1 hour at RT. Next, the sections were incubated overnight at 4°C with primary antibody (prepared in blocking buffer) against KRT5 (Basal cell marker, 2 μg/ml, Cat# PA1-37974, Thermo Scientific). The next day, slides were washed three times with PBS and incubated with fluorescently labelled secondary antibody (2 μg/ml, Cat# A-11035, Goat anti-rabbit Alexa Fluor 546, Thermo Scientific) for 1 hour at RT. The sections were mounted using VECTASHIELD mounting media (Cat# H-1200-10, Vector Laboratories, Burlingame, CA, USA) with DAPI (to stain nuclei).

For IF top staining, the cells were permeabilized with 0.1% Triton-X 100 (Cat# 194854, MP Biomedicals, Irvine, CA, USA), blocked with 10% goat serum, and incubated with primary antibodies against acetylated tubulin (Ciliated cell marker, 5 μg/ml, Cat# T7451, Sigma Aldrich), SCGB1A1 (Club cell marker, 5 μg/ml, Cat# RD181022220-01, BioVendor LLC), MUC5AC (Goblet cell marker, 1.4 μg/ml, Cat# MA5-12178, Thermo Scientific) and NOTCH3 (10 μg/ml, Cat# PA5-13203, Thermo Scientific) for 2 hours at RT. Next, the cells were washed three times with PBS and incubated with fluorescently labelled secondary antibodies (2 μg/ml, Goat anti-mouse Alexa Fluor 488, Cat# A11029; Goat anti-rabbit Alexa Fluor 546, Cat# A11035, Thermo Scientific) for 1 hour at RT. Nuclei were stained with DAPI. Finally, the ALI membranes were cut out from the Transwell® insert and mounted on a glass slide using the ProLong™ Gold Antifade mounting media without DAPI (Cat# P36930, Thermo Scientific, Waltham). Images were taken using an Olympus BX43 upright fluorescent microscope (Olympus Corporation). Confocal microscopy of NOTCH3 was performed using a Zeiss LSM 780 Confocal microscope (Carl Zeiss LLC, White Plains, NY, USA). To quantify the IF microscopy data from ALI cross-sections, at least 10 images across the whole section were captured and a minimum of 500 cells counted per sample. For quantification of top-staining, at least 4 random images of the epithelium were taken and a minimum of 1000 cells counted per sample. The number of cells positive for each marker of interest was counted using the ImageJ software (version 1.8.0_112, NIH), and normalized to the number of nuclei.

#### RNA sequencing

Genome-wide transcriptome changes in response to CSE treatment were assessed by RNA sequencing in both nonsmoker (n=4) and COPD smoker (n=4) *in vitro* ALI differentiated airway epithelium. Cells were either untreated or treated with 2.5% CSE for 7 days (ALI day 28-35) and total RNA extracted from each sample (described above). RNA-sequencing was performed on a NextSeq 500 Flowcell, High SR75 (Illumina, San Diego, CA, USA) following library preparation using the QuantSeq 3′ mRNA-Seq Library Prep Kit FWD for Illumina (Lexogen, Vienna, Austria). The library preparation and sequencing were performed by the Clinical Genomics Core at the Oklahoma Medical Research Foundation. The raw data are publicly available at the Gene Expression Omnibus (GEO) site (http://www.ncbi.nlm.nih.gov/geo/), accession number GSE152446. RNA-seq data processing followed the guidelines and practices of the ENCODE and modENCODE consortia regarding proper experimental replication, sequencing depth, data and metadata reporting, and data quality assessment (https://www.encodeproject.org/documents/cede0cbe-d324-4ce7-ace4-f0c3eddf5972/). Raw sequencing reads (in a FASTQ format) were trimmed of residual adaptor sequences using Scythe software. Low quality bases at the beginning or the end of sequencing reads were removed using sickle then the quality of remaining reads was confirmed with FastQC. Further processing of quality sequencing reads was performed with utilities provided by the Tuxedo Suite software. Reads were aligned to the Homo sapiens genome reference (GRCh38/hg38) using the TopHat component, then cuffquant and cuffdiff were utilized for gene-level read counting and differentially expression analysis. Genes that were differentially expressed in response to CSE treatment were determined using a threshold of p<0.05 on the false discovery rate (FDR). Ingenuity Pathway Analysis (IPA) (Qiagen, Redwood City, CA, USA) was used to identify molecular pathways altered in response to CSE using an unrestricted analysis.

#### Western blotting

Total protein was extracted by directly lysing the cells in the ALI wells using 100 μl of RIPA buffer (Cat# R0278, Sigma Aldrich) with 1X protease inhibitor (Cat# 88666, Thermo Scientific), 1X phosphatase inhibitor (Cat# 1862495, Thermo Scientific, Waltham), 50 mM DTT (Cat# 646563, Sigma Aldrich), and 1X NuPAGE LDS sample buffer (NP0007, Thermo Scientific). The protein lysate was boiled for 10 minutes and run on a NuPAGE 4-12% Bis-Tris gradient gel (Thermo Scientific) using the NuPAGE™ MES SDS Running buffer (NP0002, Thermo Scientific). The gel was transferred onto a nitrocellulose membrane (Amersham™ Protran™, GE Healthcare Life Sciences, Marlborough, MA, USA), using the Bio-Rad semidry transfer apparatus before western analysis. The membranes were then blocked overnight at 4°C in 4% blocking reagent (non-fat milk) made in PBS containing 0.1% Tween-20 (PBST). After blocking the membranes overnight, immobilized proteins were reacted with the following primary antibodies in 4% blocking reagent for 1 hr, at room temperature (RT) with shaking: NOTCH1 (1:3000 dilution, Cat# 4380), NOTCH2 (1:3000 dilution, Cat# 4530), NOTCH3 (1:3000 dilution, Cat# 5276) (Cell Signaling Technologies, Danvers, MA, USA), and β-tubulin (1:5000 dilution, Cat# PA5-16863, Thermo Scientific). Next, the membranes were washed 3x for 5 min each with PBST, followed by incubation with an anti-rabbit (Cat# 31462, Thermo Scientific) or anti-mouse secondary antibody (Cat# 31432, Thermo Scientific) conjugated to horseradish peroxidase (prepared in 4% blocking reagent) for 1 hr at RT with shaking. On completion of the secondary antibody incubation, the membranes were then washed again three times for 5 min with PBST and twice with PBS, and antibodies visualized after the addition of ECL reagent (Cat# 32106, Thermo Scientific) and exposure to X-ray film. The abundance of the NOTCH1-3 proteins (relative to β-tubulin levels) was quantified using the ImageJ software (version 1.8.0_112, NIH).

#### Lentivirus-based overexpression of NICD3

Generation of control or NICD3 expressing replication deficient lentiviruses was described previously (E1). Stocks of Lenti-Control and Lenti-NICD3 were titrated by qPCR (Cat#: LV900, ABM^®^good, Richmond, BC, Canada) and their infectivity confirmed by flow cytometry. To ensure >90% of cells were infected (GFP+) with each virus, HBECs were infected at a multiplicity of infection (MOI) of 50 (5 × 10^6^ viral genomes/1 × 10^5^ cells) at the time of seeding the cells on ALI. To aid virus infection, the BLEAM media was supplemented with 2 μg/ml of polybrene (Cat#: TR-1003-G, Sigma Aldrich). The next day, fresh media was added to the wells and the standard ALI protocol was continued until the day of harvest (ALI day 7).

#### siRNA mediated knockdown of NOTCH3

Primary HBECs were either transfected with 1 pmol of control siRNA (Cat# 4390844, Thermo Scientific) or NOTCH3 siRNA (Cat # 4392420, Thermo Scientific, Waltham) using Lipofectamine RNAiMax Reagent (Cat# 13778-045, Thermo Scientific) and OptiMEM media (Cat# 1985-010, Thermo Scientific) at the time of seeding the cells in ALI culture. The next day, media in both apical and basolateral chambers was replaced with fresh BLEAM media, and the standard ALI protocol was continued until day of harvest (ALI day 7).

#### Statistics

Statistical analysis was performed by two-tailed Mann Whitney U test using the GraphPad Prism version 8.0. A p value of ≤ 0.05 was considered significant.

---

**Table E1:** Demographics of primary human bronchial epithelial cell (HBEC) donors

| Phenotype | Batch number | Age (Years) | Gender | Race<sup>1</sup> | Smoker |
| --- | --- | --- | --- | --- | --- |
| **Nonsmoker** |  |  |  |  |  |
| Donor 1 | 0000501936 | 42 | Female | H | No |
| Donor 2 | 0000544414 | 48 | Male | C | No |
| Donor 3 | 0000619261 | 53 | Male | C | No |
| Donor 4 | 0000543643 | 57 | Female | C | No |
| Donor 5 | 0000613375 | 65 | Female | B | No |
| Donor 6 | 0000529235 | 67 | Female | B | No |
| Donor 7 | 0000420927 | 69 | Female | H | No |
| Donor 8 | 0000444771 | 69 | Male | B | No |
| **COPD** |  |  |  |  |  |
| Donor 1 | 0000436083 | 59 | Male | C | Yes |
| Donor 2 | 0000636518 | 62 | Female | B | Yes |
| Donor 3 | 0000440551 | 63 | Male | B | Yes |
| Donor 4 | 0000430905 | 66 | Male | C | Yes |

<sup>1</sup>Abbreviations: B = Black; C = Caucasian; H = Hispanic

**Figure E1.**
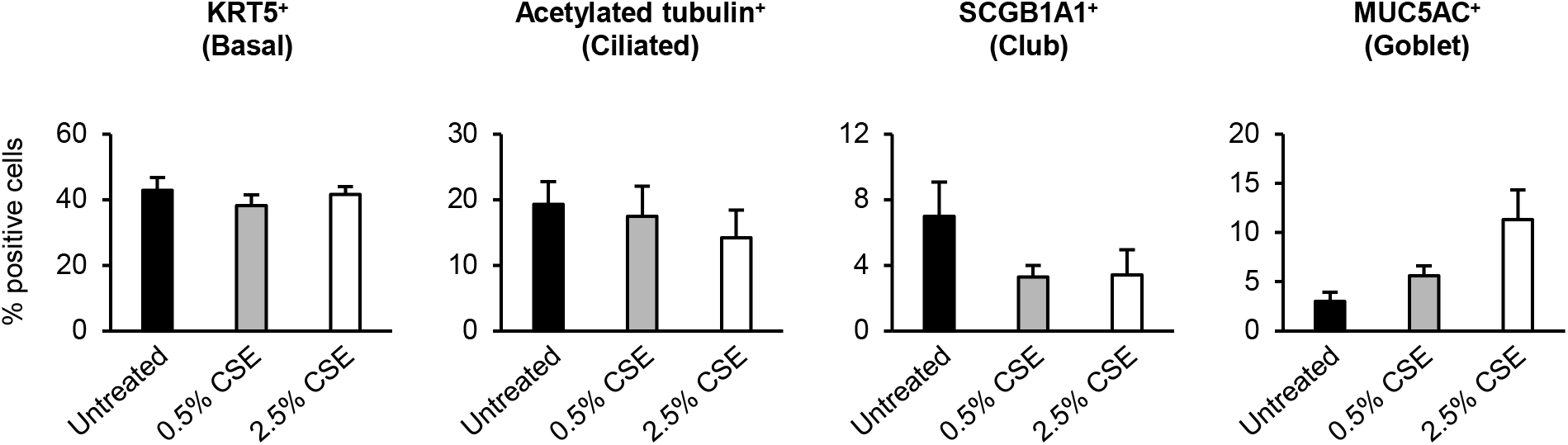
Primary human bronchial epithelial cells (HBECs) from nonsmokers were cultured on ALI for 28 days to differentiate into a pseudostratified epithelium containing basal, ciliated and secretory (club and goblet) cells. At ALI day 28, the cultures are then untreated or treated for 7 days with cigarette smoke extract (CSE). Immunofluorescent staining of basal cells (KRT5), ciliated cells (acetylated tubulin), club cells (SCGB1A1), and goblet cells (MUC5AC). Images were quantified using the ImageJ software (NIH). Data presented as mean percentage of positive cells for each condition from n=4 independent experiments. Error bars indicate SEM.

**Figure E2.**
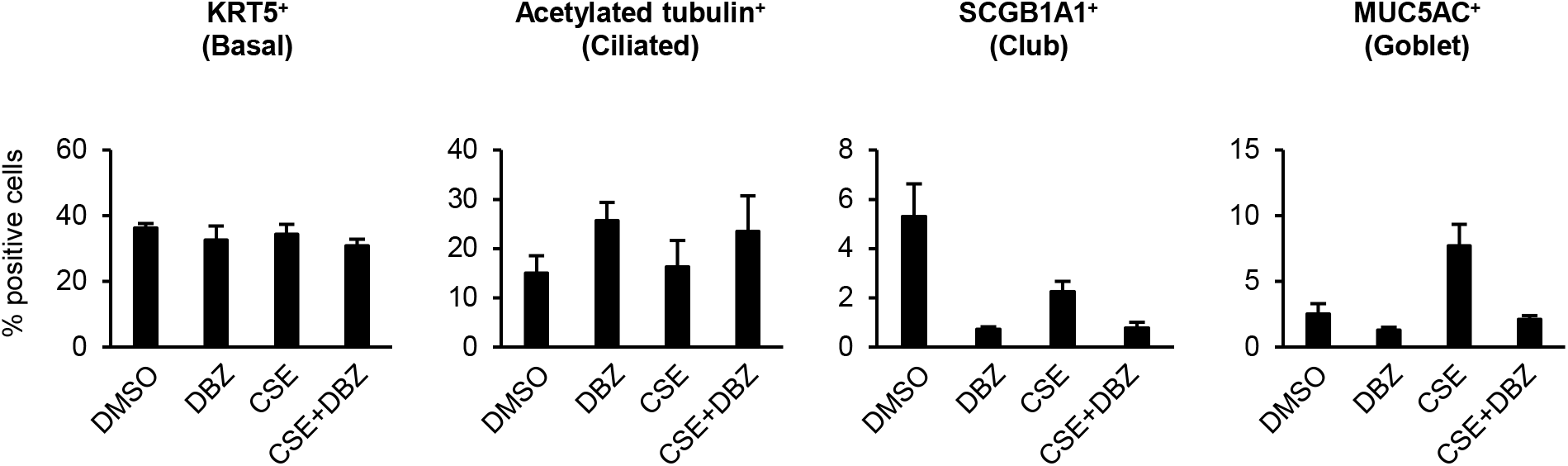
Primary human bronchial epithelial cells (HBECs) from nonsmokers were cultured on ALI for 28 days to differentiate into a pseudostratified epithelium containing basal, ciliated and secretory (club and goblet) cells. At ALI day 28, the cultures were treated with DMSO (vehicle), DBZ (0.1 μM), DMSO + CSE (2.5%) and CSE + DBZ for 7 days. Immunofluorescent staining of basal cells (KRT5), ciliated cells (acetylated tubulin), club cells (SCGB1A1), and goblet cells (MUC5AC). Images were quantified using the ImageJ software (NIH). Data presented as mean percentage of positive cells for each condition from n=4 independent experiments. Error bars indicate SEM.

**Figure E3.**
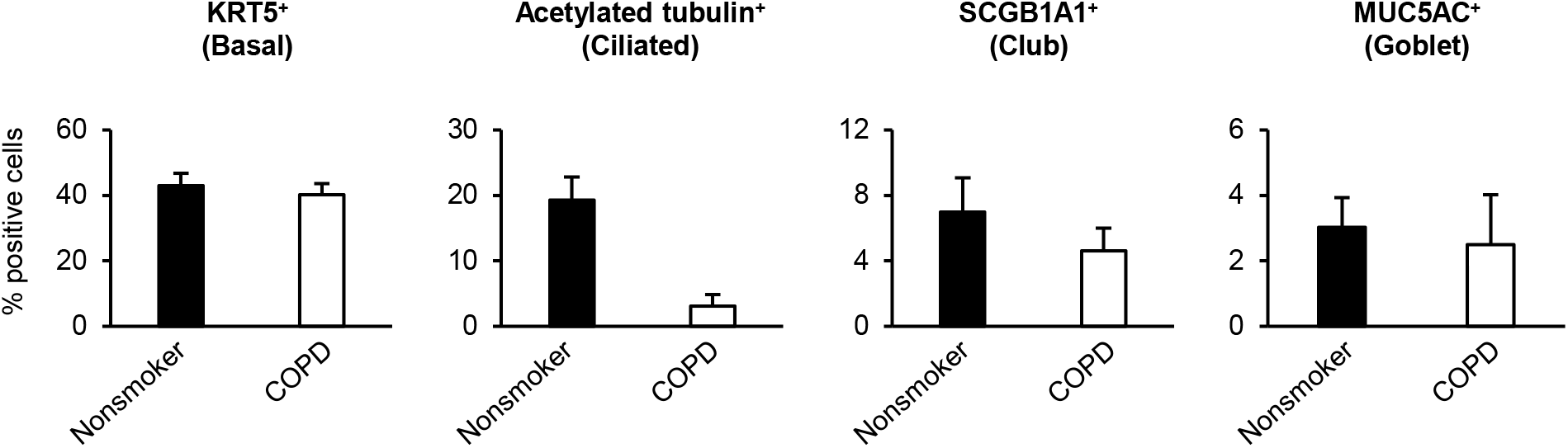
Primary human bronchial epithelial cells (HBECs) from nonsmokers and COPD smokers were cultured on ALI for 35 days to differentiate into a pseudostratified epithelium containing basal, ciliated and secretory (club and goblet) cells. Immunofluorescent staining of basal cells (KRT5), ciliated cells (acetylated tubulin), club cells (SCGB1A1), and goblet cells (MUC5AC). Images were quantified using the ImageJ software (NIH). Data presented as mean percentage of positive cells for each phenotype from n=4 independent experiments. Error bars indicate SEM.

**Figure E4.**
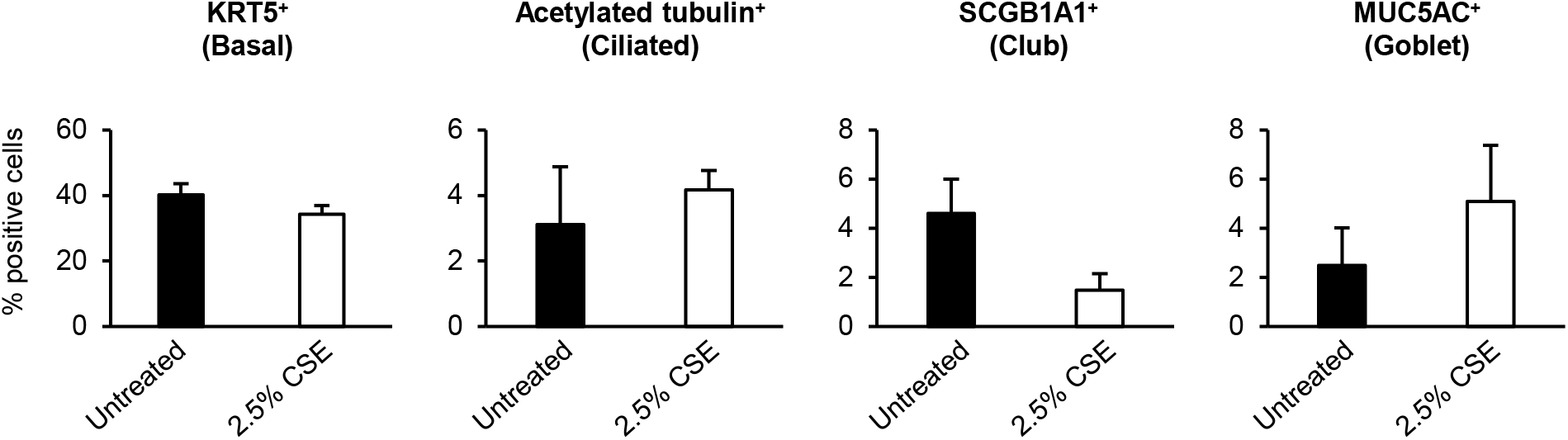
Primary human bronchial epithelial cells (HBECs) from COPD smokers were cultured on ALI for 28 days to differentiate into a pseudostratified epithelium containing basal, ciliated and secretory (club and goblet) cells. At ALI day 28, the cultures were then untreated or treated for 7 days with 2.5% cigarette smoke extract (CSE) before harvest. Immunofluorescent staining of basal cells (KRT5), ciliated cells (acetylated tubulin), club cells (SCGB1A1), and goblet cells (MUC5AC). Images were quantified using the ImageJ software (NIH). Data presented as mean percentage of positive cells for each condition from n=4 independent experiments. Error bars indicate SEM.

**Figure E5.**
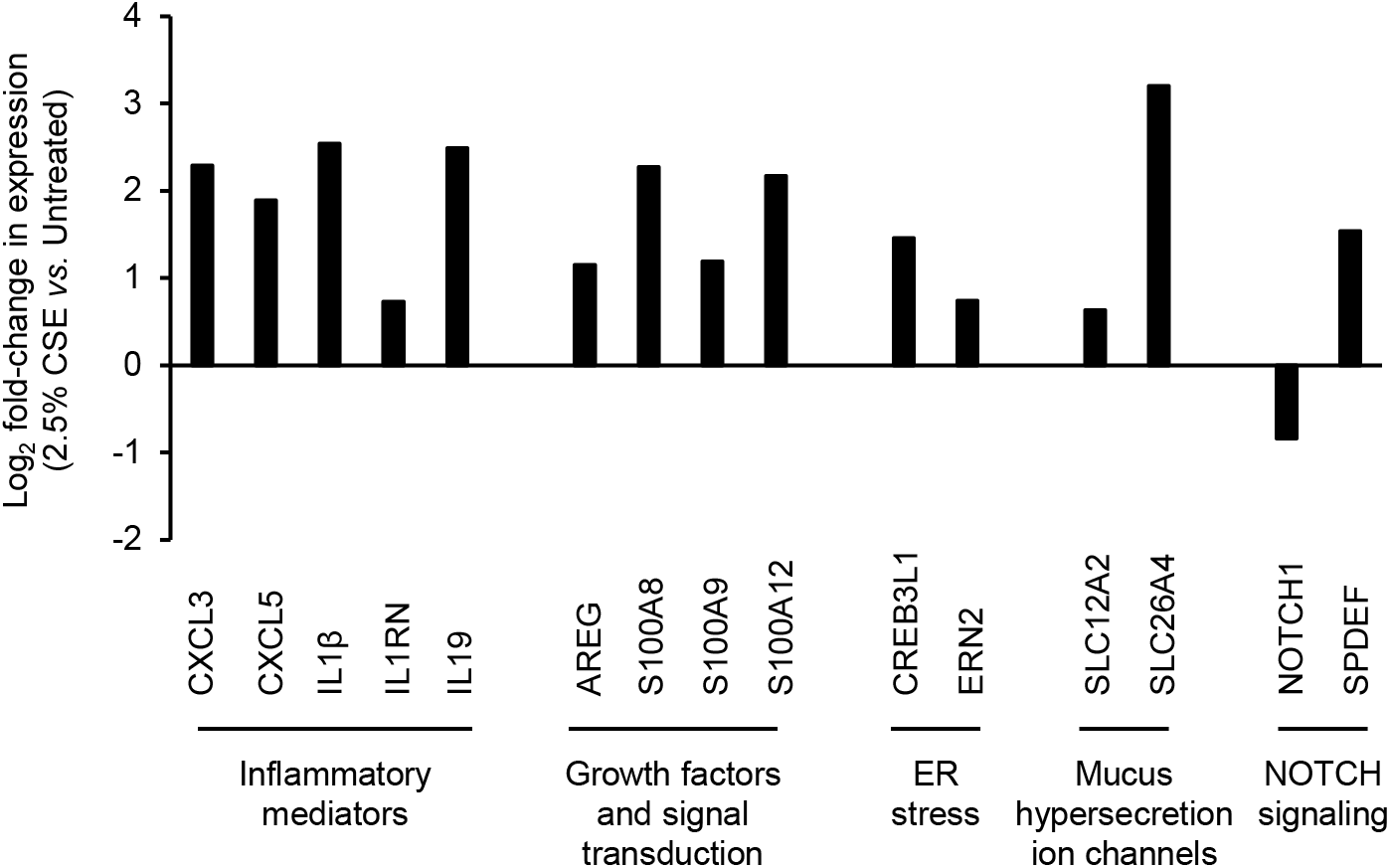
Primary human bronchial epithelial cells (HBECs) from COPD smokers were cultured on ALI for 28 days to differentiate into a pseudostratified epithelium containing basal, ciliated and secretory (club and goblet) cells. At ALI day 28, the cultures were then untreated or treated for 7 days with 2.5% cigarette smoke extract (CSE) before harvest for RNA-Seq analysis. Expression changes in genes associated with goblet cell biology present in the 230 CSE-responsive genes and also altered by CSE exposure of nonsmoker airway epithelium. Data presented as mean Log_2_ fold-change in expression (2.5% CSE treated vs. Untreated) from n=4 independent experiments.

## Notes

**Funding:** This work was supported by the following grants awarded to MSW: National Institute of General Medical Sciences (NIGMS) COBRE (GM103636, Project 4), Oklahoma Center for Adult Stem Cell (OCASCR) Research Grant, Oklahoma Shared Clinical & Translational Resources (OSCTR) Pilot Project Grant (U54GM104938), College of Medicine Alumni Association (COMAA) Research Grant, Presbyterian Health Foundation (PHF) 3D Bio-Printing Seed Grant and PHF New Investigator Seed Grant. The funders had no role in study design, data collection and analysis, decision to publish, or preparation of the manuscript.

### Competing Interest Statement

The authors have declared no competing interest.

## References

1. Tam A, Sin DD. Pathobiologic mechanisms of chronic obstructive pulmonary disease. Med Clin North Am 2012;96(4):681–698.

2. Higham A, Quinn AM, Cancado JED, Singh D. The pathology of small airways disease in copd: Historical aspects and future directions. Respir Res 2019;20(1):49.

3. Lopez-Campos JL, Tan W, Soriano JB. Global burden of copd. Respirology 2016;21(1):14–23.

4. Laniado-Laborin R. Smoking and chronic obstructive pulmonary disease (copd). Parallel epidemics of the 21 century. Int J Environ Res Public Health 2009;6(1):209–224.

5. Hogan BL, Barkauskas CE, Chapman HA, Epstein JA, Jain R, Hsia CC, Niklason L, Calle E, Le A, Randell SH, et al. Repair and regeneration of the respiratory system: Complexity, plasticity, and mechanisms of lung stem cell function. Cell Stem Cell 2014;15(2):123–138.

6. Whitsett JA, Kalin TV, Xu Y, Kalinichenko VV. Building and regenerating the lung cell by cell. Physiol Rev 2019;99(1):513–554.

7. Tilley AE, Walters MS, Shaykhiev R, Crystal RG. Cilia dysfunction in lung disease. Annu Rev Physiol 2015;77:379–406.

8. Benam KH, Vladar EK, Janssen WJ, Evans CM. Mucociliary defense: Emerging cellular, molecular, and animal models. Ann Am Thorac Soc 2018;15(Suppl 3):S210–S215.

9. Whitsett JA. Airway epithelial differentiation and mucociliary clearance. Ann Am Thorac Soc 2018;15(Suppl 3):S143–S148.

10. Reid AT, Veerati PC, Gosens R, Bartlett NW, Wark PA, Grainge CL, Stick SM, Kicic A, Moheimani F, Hansbro PM, et al. Persistent induction of goblet cell differentiation in the airways: Therapeutic approaches. Pharmacol Ther 2018;185:155–169.

11. Kiyokawa H, Morimoto M. Notch signaling in the mammalian respiratory system, specifically the trachea and lungs, in development, homeostasis, regeneration, and disease. Dev Growth Differ 2020;62(1):67–79.

12. Reid AT, Nichol KS, Chander Veerati P, Moheimani F, Kicic A, Stick SM, Bartlett NW, Grainge CL, Wark PAB, Hansbro PM, et al. Blocking notch3 signaling abolishes muc5ac production in airway epithelial cells from individuals with asthma. Am J Respir Cell Mol Biol 2020;62(4):513–523.

13. Boucherat O, Chakir J, Jeannotte L. The loss of hoxa5 function promotes notch-dependent goblet cell metaplasia in lung airways. Biol Open 2012;1(7):677–691.

14. Gomi K, Arbelaez V, Crystal RG, Walters MS. Activation of notch1 or notch3 signaling skews human airway basal cell differentiation toward a secretory pathway. PLoS One 2015;10(2):e0116507.

15. Danahay H, Pessotti AD, Coote J, Montgomery BE, Xia D, Wilson A, Yang H, Wang Z, Bevan L, Thomas C, et al. Notch2 is required for inflammatory cytokine-driven goblet cell metaplasia in the lung. Cell Rep 2015;10(2):239–252.

16. Tsao PN, Wei SC, Wu MF, Huang MT, Lin HY, Lee MC, Lin KM, Wang IJ, Kaartinen V, Yang LT, et al. Notch signaling prevents mucous metaplasia in mouse conducting airways during postnatal development. Development 2011;138(16):3533–3543.

17. Jing Y, Gimenes JA, Mishra R, Pham D, Comstock AT, Yu D, Sajjan U. Notch3 contributes to rhinovirus-induced goblet cell hyperplasia in copd airway epithelial cells. Thorax 2019;74(1):18–32.

18. Carrer M, Crosby JR, Sun G, Zhao C, Damle SS, Kuntz SG, Monia BP, Hart CE, Grossman TR. Antisense oligonucleotides targeting jagged 1 reduce house dust mite-induced goblet cell metaplasia in the adult murine lung. Am J Respir Cell Mol Biol 2020.

19. Lafkas D, Shelton A, Chiu C, de Leon Boenig G, Chen Y, Stawicki SS, Siltanen C, Reichelt M, Zhou M, Wu X, et al. Therapeutic antibodies reveal notch control of transdifferentiation in the adult lung. Nature 2015;528(7580):127–131.

20. Rock JR, Gao X, Xue Y, Randell SH, Kong YY, Hogan BL. Notch-dependent differentiation of adult airway basal stem cells. Cell Stem Cell 2011;8(6):639–648.

21. Guseh JS, Bores SA, Stanger BZ, Zhou Q, Anderson WJ, Melton DA, Rajagopal J. Notch signaling promotes airway mucous metaplasia and inhibits alveolar development. Development 2009;136(10):1751–1759.

22. Gomi K, Staudt MR, Salit J, Kaner RJ, Heldrich J, Rogalski AM, Arbelaez V, Crystal RG, Walters MS. Jag1-mediated notch signaling regulates secretory cell differentiation of the human airway epithelium. Stem Cell Rev Rep 2016;12(4):454–463.

23. Siebel C, Lendahl U. Notch signaling in development, tissue homeostasis, and disease. Physiol Rev 2017;97(4):1235–1294.

24. Steiling K, van den Berge M, Hijazi K, Florido R, Campbell J, Liu G, Xiao J, Zhang X, Duclos G, Drizik E, et al. A dynamic bronchial airway gene expression signature of chronic obstructive pulmonary disease and lung function impairment. Am J Respir Crit Care Med 2013;187(9):933–942.

25. Tilley AE, O’Connor TP, Hackett NR, Strulovici-Barel Y, Salit J, Amoroso N, Zhou XK, Raman T, Omberg L, Clark A, et al. Biologic phenotyping of the human small airway epithelial response to cigarette smoking. PLoS One 2011;6(7):e22798.

26. Fujisawa T, Velichko S, Thai P, Hung LY, Huang F, Wu R. Regulation of airway muc5ac expression by il-1beta and il-17a; the nf-kappab paradigm. J Immunol 2009;183(10):6236–6243.

27. Zuo WL, Yang J, Gomi K, Chao I, Crystal RG, Shaykhiev R. Egf-amphiregulin interplay in airway stem/progenitor cells links the pathogenesis of smoking-induced lesions in the human airway epithelium. Stem Cells 2017;35(3):824–837.

28. Wang G, Xu Z, Wang R, Al-Hijji M, Salit J, Strulovici-Barel Y, Tilley AE, Mezey JG, Crystal RG. Genes associated with muc5ac expression in small airway epithelium of human smokers and non-smokers. BMC Med Genomics 2012;5:21.

29. Kang JH, Hwang SM, Chung IY. S100a8, s100a9 and s100a12 activate airway epithelial cells to produce muc5ac via extracellular signal-regulated kinase and nuclear factor-kappab pathways. Immunology 2015;144(1):79–90.

30. Park KS, Korfhagen TR, Bruno MD, Kitzmiller JA, Wan H, Wert SE, Khurana Hershey GK, Chen G, Whitsett JA. Spdef regulates goblet cell hyperplasia in the airway epithelium. J Clin Invest 2007;117(4):978–988.

31. Marcet B, Chevalier B, Luxardi G, Coraux C, Zaragosi LE, Cibois M, Robbe-Sermesant K, Jolly T, Cardinaud B, Moreilhon C, et al. Control of vertebrate multiciliogenesis by mir-449 through direct repression of the delta/notch pathway. Nat Cell Biol 2011;13(6):693–699.

32. Mori M, Mahoney JE, Stupnikov MR, Paez-Cortez JR, Szymaniak AD, Varelas X, Herrick DB, Schwob J, Zhang H, Cardoso WV. Notch3-jagged signaling controls the pool of undifferentiated airway progenitors. Development 2015;142(2):258–267.

33. Paul MK, Bisht B, Darmawan DO, Chiou R, Ha VL, Wallace WD, Chon AT, Hegab AE, Grogan T, Elashoff DA, et al. Dynamic changes in intracellular ros levels regulate airway basal stem cell homeostasis through nrf2-dependent notch signaling. Cell Stem Cell 2014;15(2):199–214.

34. Zhang S, Loch AJ, Radtke F, Egan SE, Xu K. Jagged1 is the major regulator of notch-dependent cell fate in proximal airways. Dev Dyn 2013;242(6):678–686.

35. Pardo-Saganta A, Law BM, Tata PR, Villoria J, Saez B, Mou H, Zhao R, Rajagopal J. Injury induces direct lineage segregation of functionally distinct airway basal stem/progenitor cell subpopulations. Cell Stem Cell 2015;16(2):184–197.

36. Tilley AE, Harvey BG, Heguy A, Hackett NR, Wang R, O’Connor TP, Crystal RG. Down-regulation of the notch pathway in human airway epithelium in association with smoking and chronic obstructive pulmonary disease. Am J Respir Crit Care Med 2009;179(6):457–466.

37. Buro-Auriemma LJ, Salit J, Hackett NR, Walters MS, Strulovici-Barel Y, Staudt MR, Fuller J, Mahmoud M, Stevenson CS, Hilton H, et al. Cigarette smoking induces small airway epithelial epigenetic changes with corresponding modulation of gene expression. Hum Mol Genet 2013;22(23):4726–4738.

38. Staudt MR, Buro-Auriemma LJ, Walters MS, Salit J, Vincent T, Shaykhiev R, Mezey JG, Tilley AE, Kaner RJ, Ho MW, et al. Airway basal stem/progenitor cells have diminished capacity to regenerate airway epithelium in chronic obstructive pulmonary disease. Am J Respir Crit Care Med 2014;190(8):955–958.

39. Ghosh M, Miller YE, Nakachi I, Kwon JB, Baron AE, Brantley AE, Merrick DT, Franklin WA, Keith RL, Vandivier RW. Exhaustion of airway basal progenitor cells in early and established chronic obstructive pulmonary disease. Am J Respir Crit Care Med 2018;197(7):885–896.

40. Stupnikov MR, Yang Y, Mori M, Lu J, Cardoso WV. Jagged and delta-like ligands control distinct events during airway progenitor cell differentiation. Elife 2019;8.

41. Ruiz Garcia S, Deprez M, Lebrigand K, Cavard A, Paquet A, Arguel MJ, Magnone V, Truchi M, Caballero I, Leroy S, et al. Novel dynamics of human mucociliary differentiation revealed by single-cell rna sequencing of nasal epithelial cultures. Development 2019;146(20).

42. Cheng Z, Tan Q, Tan W, Zhang LI. Cigarette smoke induces the expression of notch3, not notch1, protein in lung adenocarcinoma. Oncol Lett 2015;10(2):641–646.

43. Li S, Hu X, Wang Z, Wu M, Zhang J. Different profiles of notch signaling in cigarette smoke-induced pulmonary emphysema and bleomycin-induced pulmonary fibrosis. Inflamm Res 2015;64(5):363–371.

44. Jung JG, Stoeck A, Guan B, Wu RC, Zhu H, Blackshaw S, Shih Ie M, Wang TL. Notch3 interactome analysis identified wwp2 as a negative regulator of notch3 signaling in ovarian cancer. PLoS Genet 2014;10(10):e1004751.

45. Jia L, Yu G, Zhang Y, Wang MM. Lysosome-dependent degradation of notch3. Int J Biochem Cell Biol 2009;41(12):2594–2598.

46. Zhang X, Wang YN, Zhu JJ, Liu XX, You H, Gong MY, Zou M, Cheng WH, Zhu JH. N-acetylcysteine negatively regulates notch3 and its malignant signaling. Oncotarget 2016;7(21):30855–30866.

47. Barnes PJ. Oxidative stress-based therapeutics in copd. Redox Biol 2020:101544.

48. Vij N, Chandramani-Shivalingappa P, Van Westphal C, Hole R, Bodas M. Cigarette smoke-induced autophagy impairment accelerates lung aging, copd-emphysema exacerbations and pathogenesis. Am J Physiol Cell Physiol 2018;314(1):C73–C87.

49. Yi G, Liang M, Li M, Fang X, Liu J, Lai Y, Chen J, Yao W, Feng X, Hu, et al. A large lung gene expression study identifying il1b as a novel player in airway inflammation in copd airway epithelial cells. Inflamm Res 2018;67(6):539–551.

50. Chen G, Sun L, Kato T, Okuda K, Martino MB, Abzhanova A, Lin JM, Gilmore RC, Batson BD, O’Neal YK, et al. Il-1beta dominates the promucin secretory cytokine profile in cystic fibrosis. J Clin Invest 2019;129(10):4433–4450.

## References

E1. Gomi K, Arbelaez V, Crystal RG, Walters MS. Activation of notch1 or notch3 signaling skews human airway basal cell differentiation toward a secretory pathway. PLoS One 2015;10(2):e0116507.

